# Copper supplementation mitigates Parkinson-like wild-type SOD1 pathology and nigrostriatal degeneration in a novel mouse model

**DOI:** 10.1101/2025.02.26.640461

**Authors:** Benjamin D. Rowlands, Benjamin G. Trist, Conor Karozis, Greta Schaffer, David Mor, Richard Harwood, Sarah A. Rosolen, Veronica Cottam, Freyja Persson-Carboni, Miriam Richardson, Anne A. Li, Michael P. Gotsbacher, Amr H. Abdeen, Rachel Codd, Kay L. Double

**Affiliations:** Brain and Mind Centre, The University of Sydney, Sydney, New South Wales 2006, Australia; School of Medical Sciences (Neuroscience), Faculty of Medicine and Health, The University of Sydney, Sydney, New South Wales 2006, Australia; Sydney Microscopy and Microanalysis, The University of Sydney, Sydney, New South Wales 2006, Australia

**Keywords:** Neurodegeneration, Superoxide dismutase 1, post-translational modification, protein misfolding, Parkinson disease, CuATSM, copper supplementation, mouse model, substantia nigra pars compacta

## Abstract

Wild-type superoxide dismutase 1 (disSOD1) protein misfolding and deposition is implicated in the death of substantia nigra (SN) dopamine neurons in Parkinson disease. Regionally reduced copper availability, and subsequent reduced copper binding to SOD1, is a key factor driving the development of this pathology, suggesting brain copper supplementation may constitute an effective means of preventing its formation. We evaluated the potential of the blood-brain-barrier-permeable copper delivery drug, CuATSM, to attenuate the misfolding and deposition of wild-type disSOD1, and associated neuron death, in a novel mouse model that expresses this pathology. Using proteomic and elemental mass spectrometry, together with biochemical and histological workflows, we demonstrated copper supplementation corrects altered post-translational modifications on soluble SOD1 and improves the enzymatic activity of the protein in the brains of these animals. These changes were associated with a significant reduction in disSOD1 pathology and preservation of dopamine neurons in the SN, which were highly correlated with tissue copper levels. These data position wild-type disSOD1 pathology as a novel drug target for Parkinson disease and suggest that brain copper supplementation may constitute an effective means of slowing SN dopamine neuron death in this disorder.

## Background

Neurological disorders are the leading cause of disability worldwide, with none growing at a faster rate than Parkinson disease^1–3^. The progressive death of dopamine-producing neurons in the substantia nigra (SN), a brain region critical for movement control, is the principal cause of disability in this disorder^4, 5^. Health systems worldwide are ill-prepared to deal with the growing population of Parkinson disease patients^6^, as there are currently no treatments that can rescue the loss of these cells. A treatment that slows the rate of cell death is consequently a primary goal of research into Parkinson disease therapeutics^6^. Putative disease-modifying treatments based upon traditional pathophysiological aspects of Parkinson disease^7^ have failed to significantly slow disease progression in clinical trials^8^, suggesting the exploration of new disease targets and therapeutic approaches is warranted.

Our group has established that the normally protective metalloprotein superoxide dismutase 1 (SOD1) is structurally abnormal and dysfunctional specifically within vulnerable brain regions in Parkinson disease, including the SN. This begins in the early stages of the disease and results in significant deposition of the protein, which is thought to contribute to the death of SN dopamine neurons^9, 10^. Unlike well-documented mutant SOD1 pathology underlying neuron death in inherited forms of amyotrophic lateral sclerosis (fALS)^11^, these alterations are not driven by *SOD1* gene mutations, but rather non-genetic alterations to wild-type SOD1^10^. Given neither abnormal SOD1 nor dopamine neuron death, can be quantified in the living human brain, we developed the first mouse model expressing Parkinson-like wild-type SOD1 pathology to better understand the relationship between this pathology and dopamine neuron vulnerability. This was achieved by replicating two key changes identified in the post-mortem Parkinson disease SN^9, 12^: elevated human SOD1 protein levels and a decrease in cellular copper levels. These **S**OD1-**O**verexpressing-**C**tr1 **K**nockdown (SOCK) mice develop abundant human wild-type SOD1 pathology from 6 weeks of age in the SN, which is followed by age-associated motor impairments and significant SN dopamine neuron loss^13^. These findings support an association between wild-type SOD1 pathology and SN dopamine neuron death and highlight the potential of SOD1 pathology as a novel disease-modifying target for Parkinson disease.

Our analyses of mechanisms driving the formation of SOD1 pathology in SOCK mice implicate a series of altered post-translational modifications^13^. Among these is a decrease in copper binding to SOD1^13^, a finding also characterizing multiple forms of mutant SOD1 protein associated with spinal motor neuron death in transgenic mouse models of fALS^14, 15^. These data suggest that Parkinson-like wild-type SOD1 pathology may be mitigated by improving soluble SOD1 copper binding.

Here we show that treatment of SOCK mice with the blood-brain-barrier-permeable copper delivery drug, diacetyl-bis(4-methylthiosemicarbazonato)copper(II) (CuATSM), mitigates the formation of Parkinson-like wild-type SOD1 pathology, increases SN dopamine neuron survival and improves motor function. These benefits are associated with normalization of the structure and enzymatic activity of the soluble protein. Our findings highlight the potential of copper supplementation approaches to confer neuroprotection in Parkinson disease.

## Methods

### CuATSM preparation

The synthesis of diacetyl-bis(N4-methyl-3-thiosemicarbazonato) copper(II) (CuATSM; CAS# 68341-09-3) was adapted from literature^16, 17^. The Australian Therapeutic Goods Administration (TGA) currently does not provide guidelines on minimum purity standards for active pharmaceutical ingredients (API) tested in pre-clinical studies. As such, CuATSM was prepared taking standards of Good Laboratory Practice (GLP) into consideration, as defined by the Food and Drug Administration (FDA) in the United States of America (Code of Federal Regulations, Title 21, Chapter I, Subchapter A, Part 58.1 – 58.219; Good Laboratory Practice for Nonclinical Laboratory Studies).

#### General

Chemicals were purchased from Sigma Aldrich (Castle Hill, NSW, AU) and used without any further purification unless stated otherwise. Ultrapure water (prepared by purifying deionized water, to attain a sensitivity of 18.2 MΩ-cm at 25 °C) was used whenever water was required.

#### Instrumentation

Nuclear magnetic resonance (NMR) spectroscopy (^1^H and ^13^C) spectra were recorded in 5-mm Pyrex tubes (Wilmad, USA) at 298 K on a Bruker Avance III 600 NMR spectrometer, fitted with a cryogenic TCI probe head and using standard pulse sequences from the Bruker library. The spectra were processed using TOPSPIN3 (Bruker, Karlsruhe, Germany). The spectral data are reported in ppm (δ) and referenced to residual solvent (deuterated dimethyl sulfoxide [DMSO-*d*_6_] 2.50/39.52 ppm). Residual water generates a signal at 3.30 ppm.

Compound purity was determined by High Pressure Liquid Chromatography Mass Spectrometry (HPLC-MS) using an Agilent system (Santa Clara, CA, USA) consisting of a 1260 series quaternary pump with an inbuilt degasser, 1200 series autosampler, temperature-controlled column compartment, diode array detector, and fraction collector, a 6120 series single quadrupole mass spectrometer and an Agilent Zorbax Eclipse XDB-C18 column (150 × 4.6 mm i.d., 5 µm) at 30 °C. The drying gas flow, temperature, and nebuliser were set to 12 L min-1, 350 °C, and 35 psi, respectively. Agilent OpenLAB Chromatography Data System (CDS) ChemStation Edition (C.01.05) was used for data acquisition and processing. Electrospray ionisation (ESI) was used to analyse aliquots (10 µL) in positive ion mode with a 3500 V capillary voltage.

#### Synthesis of Diacetyl-bis(N4-methyl-3-thiosemicarbazone) (ATSMH_2_; CAS#63618-91-7)

4-methyl-3-thiosemicarbazide (H_2_NNHCSNHCH_3_) (6.6 g, 63 mmol) was dissolved in 125 mL of distilled water containing 5% acetic acid at 60 °C. The solution was cooled to 10 °C and to this was added 6.12 mL (70 mmol) of 2,3-butanedione dropwise over 20 min. The reaction was allowed to stir overnight at room temperature. After cooling the solution in an ice-bath to 4 °C, the precipitate was retrieved by filtration and washed with 3×100 mL of distilled water. The precipitate was dissociated into fine powder and washed with 2×30 mL of ice-cold diethyl ether, then dried *in vacuo* to yield ATSMH_2_ as off-white crystalline powder (7.81 g, 72 %). NMR (DMSO-*d*_6_, 600 MHz; **Supplementary Figure 1**) δ_H_ 10.21 (s, 2H, H-4), 8.37 (m, 2H, H-2), 3.02 (d, 6H, *J* = 4.5 Hz, H-1), 2.20 (s, 6H, H-7). δ_C_ 178.46 (C-3), 147.97 (C-6), 31.20 (C-1), 11.67 (C-7)(**Supplementary Figure 2**). MS ESI+ m/z_calc_ 261.1, observed [M+H^+^]^+^ 261.1 (100), [M+Na^+^]^+^ 283.1.

#### Synthesis of CuATSM

ATSMH_2_ (1.0 g, 3.8 mmol) was suspended in dimethylformamide (25 mL) and Cu(OAc)_2_.H_2_O (0.80 g, 4.0 mmol) was added in small portions. The reaction was stirred at reflux for 4 h and allowed to cool to room temperature. The red-brown precipitate was filtered, rinsed with 10 mL cold methanol and 2×30 mL of ice-cold diethyl ether, and dried *in vacuo* to give Cu(II)ATSM as a brown powder (1.1 g, 85%). High-resolution MS (ESI+): expected for C_8_H_15_N_6_S_2_Cu^+^ *m/z* 322.00902/324.00721 as [M+H^+^]^+^ for [63/65]Cu, found *m/z* 322.00876/324.00620 (+0.79/+3.11 ppm)(**Supplementary Figures 3-4**). The chemical purity of CuATSM was validated at >90%.

### Animals

All methods conformed to the Australian Code of Practice for the Care and Use of Animals for Scientific Purposes^18^, with protocols approved by the Animal Ethics Committee at the University of Sydney (Ethics ID: 2020_1849). Hemizygous male mice expressing seven-transgene copies of the human *SOD1^WT^* mouse strain (B6.Cg-Tg(SOD1)2Gur/J) were crossbred with female *Ctr1^+/−^* (*Slc31a1^tm^*^2^*^.1Djt^*/J) mice to produce the novel h*SOD1^WT^*/*Ctr1^+/−^* (SOCK) mouse strain, as well as wild-type C57BL/6 littermates. Both h*SOD1*^WT^ and *Ctr1^+/−^* mouse lines were sourced from The Jackson Laboratory (Bar Harbor, Maine, USA) and all mouse lines were bred and maintained at Australian BioResources (Moss Vale, NSW, Australia). Mice were housed in individually ventilated cages (Tecniplast) (12/12 h light/dark cycle, 22 °C, 45% humidity) with nesting material, and tinted polycarbonate igloos and tubes for environmental enrichment. Enclosures contained Breeder’s Choice Cat Litter with paper tissues provided for bedding and *ad libitum* access to standard chow pellets and water. Tail snips were obtained from all mice prior to the age of weaning (3 weeks) for commercial genotyping (Transnetyx) of both *Slc31a1* and h*SOD1^WT^* genes. At 7 weeks old, mice were transported to the Laboratory Animal Services facility at the Charles Perkins Centre (University of Sydney) where they acclimated to the new environment for one week. Mice were monitored twice per week where general behaviour, appearance and weight data were collected as measures of overall health. Pilot phase (n=27) and treatment phase (n=80) group sizes were calculated *a priori* using G*Power (95% power, 2-tailed t-test, 5% significance level) from variance and effect sizes observed in brain copper levels from previous work where wildtype and *Ctr1*^+/−^ mice were treated using CuATSM^19^.

### Treatments and tissue collection

CuATSM was administered orally using the micropipette-guided drug administration method^20^ in two phases. The first was a pilot experiment conducted in 27 wild-type mice to determine the optimal concentration of CuATSM to increase brain copper levels, whereby 8-week-old mice were treated daily for three weeks before tissues were harvested and analysed for their metal contents using inductively-coupled plasma mass spectrometry (ICP-MS), as described below. This identified 15 mg/kg CuATSM as a suitable dose for the second phase of our treatment experiments, in which 10 mice of each experimental genotype (wild-type, h*SOD1*^WT^, *Ctr1^+/−^,* SOCK – equal mix of sexes) were treated daily with either CuATSM or vehicle (3 parts condensed milk to 10 parts 0.9% NaCl containing 0.5% (v/v) benzyl alcohol, 0.4% (v/v) Tween-80 and 0.5% (w/v) Na-carboxymethylcellulose) from 8 weeks of age until 5.5 months of age. Following treatment, mice were deeply anaesthetized in an induction chamber using 5% isoflurane for 1 min, then transferred to a nose cone (2% isoflurane) and perfused at 2 mL/min for 5 min with chilled phosphate-buffered saline (PBS) through the left ventricle, prior to harvesting of the brain, lumbar spinal cord and liver. Brains were bisected sagittally into two hemispheres, before regions of interest (midbrain, striatum) were micro-dissected from the left hemisphere and stored with liver tissues at −80 °C for biochemical analyses. The lumbar spinal cord was bisected coronally and the superior half frozen at −80 °C for biochemical analyses. The right brain hemisphere and inferior half of the lumbar spinal cord were placed in 4% v/v paraformaldehyde in 1x PBS overnight in preparation for immunofluorescent staining.

### Fresh mouse tissue preparation for biochemical measurements

Fresh tissues were homogenized in 5x homogenization buffer volume (µL) per mg tissue weight (20 mM Tris-base pH 7.4 containing EDTA-free protease inhibitor (Sigma-Aldrich) and phosphatase inhibitor (Roche) using a Kontes pestle pellet mechanical tissue grinder (Sigma-Aldrich). Following homogenization, extracts were incubated at 4 °C for 30 min before protein concentration was determined using a bicinchoninic acid assay according to the manufacturer’s instructions (Thermo-Fisher Scientific).

### SOD1 immunoprecipitation and preparation for proteomic mass spectrometry

SOD1 protein was immunoprecipitated from mouse brain tissue homogenates as previously described^21^. Briefly, 100 µg polyclonal SOD1 antibody (Enzo Life Sciences, Farmingdale, NY, USA; **Supplementary Table 1**) was conjugated to 10 mg Dynabeads M-280 Tosyl-activated (40 mg beads/mL; Invitrogen, Carlsbad, CA, USA) in coupling buffer (0.1 M boric acid, pH 9.5; 1.2 M ammonium sulphate) overnight at 37 °C. After being blocked with 0.5% w/v BSA in PBS (pH 7.4), Dynabeads were washed with 0.1% w/v BSA in PBS (pH 7.4) and incubated with tissue homogenates (200 µg total protein) diluted in PBS (pH 7.4) to 40 mg beads/mL overnight at 4 °C. Dynabeads were then washed with PBS (pH 7.4) and immunoprecipitated proteins eluted using 0.1 M glycine (pH 3). After being dried under pressure using a vacuum concentrator, immunoprecipitates were resuspended in 50 mM ammonium bicarbonate (pH 8) containing 6 M urea and reduced with dithiothreitol (DTT; 10 mM final) for 30 min at 56 °C, before being alkylated with iodoacetamide (IAA; 20 mM final) for 30 min at room temperature in the dark and quenched with a further 10 mM DTT for 30 min at room temperature. Samples were then diluted five-fold using 50 mM ammonium bicarbonate (pH 8), acetonitrile added to 10%, and in-solution digestion performed overnight at room temperature using 0.2 µg sequencing-grade modified trypsin (Promega, Madison, WI, USA). Samples were acidified using trifluoroacetic acid, desalted using Pierce C18 Tips (ThermoFisher Scientific, USA), dried under pressure, resuspended in loading buffer (0.1% formic acid, 3% ACN) and transferred to HPLC vials immediately prior to mass spectrometry analyses.

### Mass spectrometry data acquisition and analysis

Peptide digests were analyzed using label-free Fourier Transform Mass Spectrometry at Sydney Mass Spectrometry (Sydney, New South Wales, Australia) as previously described^13^. Briefly, analyses utilized an UltiMate 3000 RSLCnano system (ThermoFisher Scientific, USA) coupled online via a Nanospray Ion Source (ThermoFisher Scientific, USA) to a Q Exactive HF-X Hybrid Quadrupole-Orbitrap Mass Spectrometer (ThermoFisher Scientific, USA). Peptides were loaded onto an in-house packed ReproSil-Pur 120 C18-AQ analytical column (75 µm id x 40 cm, 1.9 µm particle size; Dr Maisch, Ammerbuch, Germany) regulated to 60 °C using a PRSO-V2 Sonation column oven (Sonation, Baden-Wuerttemberg, Germany). Peptide elution was performed over 90 min using a binary gradient of solvent A (0.1% formic acid in MilliQ water) to solvent B (0.1% formic acid in 80% ACN diluted with MilliQ water) at a separation flow rate of 300-450 nL/min. Data were acquired using Xcalibur software (Version 4.4.16.14, ThermoFisher Scientific, USA) in a data-independent (DIA) mode. The mass spectrometer operated in a positive ion mode at a 2.4 kV needle voltage in a data-independent fashion, with 20 dynamic DIA segments covering the mass range from 350 to 1650 *m/z*. The resolution for the MS1 scan was set to 60k with a max injection time of 50 ms and an AGC target of 3e6. The DIA scans were acquired in the Orbitrap with a resolution of 30k after fragmentation in the HCD cell (max injection time: auto; AGC target: 3e6; fixed first mass: 300 *m/z*; loop count: 1; MSX count: 1; isolation window: 26-589 *m/z*).

Raw DIA data was processed using Spectronaut software’s directDIA workflow (Version 19, Biognosys, Zurich, Switzerland). Raw data files were used to generate an *in silico* spectral library using Spectronaut Pulsar, which was performed using BGS factory settings, with the Human Uniprot fasta file employed as a protein database for searches. Two missed cleavages were allowed and the false discovery rate (FDR) controlled at 1% for both PSM and protein group levels. Peptide identification and label-free quantification in each individual sample were then performed using default BGS factory settings, with spectra screened against Uniprot entry P00441 (SODC-HUMAN), corresponding to human SOD1. Cross-run normalization was disabled, PTM localization was enabled (summative PTM consolidation strategy and a probability cutoff of 0.75), and analyses were performed with a log2 ratio candidate filter of 0.58, a confidence (Q) candidate filter of 0.05 and multiple comparisons testing correction enabled. Carbamidomethyl (C) was included as a fixed modification, while modifications of interest were included individually in variable modifications alongside acetylation (N-term) and oxidation (M). PTMs of interest were analyzed in separate analysis batches, including; acetylation (K; +42.01), acetylglucosamine (NST; 203.08) carboxymethyllysine (K; 58.01), deamidation (N/Q; +0.98), kynurenine (W; +3.99), glycation (K/R; +108.02 with neutral loss of three water molecules^22^), glycosylation (NST; +162.05), nitration (W; +44.99), oxidation (H/W; +15.99), phosphorylation (S/T; +79.97), succinylation (K; +100.02), ubiquitination (GlyGly, K; 114.04). A maximum of 5 modifications per peptide was allowed, with two missed trypsin cleavages. SOD1 protein was not identified in negative control immunoprecipitates prepared using Dynabeads that were not conjugated to our capture antibody, suggesting negligible false discovery of SOD1 protein in tissue extracts^21^. No differences in the relative levels of PTMs of interest were identified between immunoprecipitated and non-immunoprecipitated commercial SOD1 protein in a previous study^21^, indicating that our immunoprecipitation protocol does not significantly alter PTMs of interest.

### SOD1 and CCS protein quantification

Immunoblotting for SOD1 protein was performed as previously described^13^. Homogenates (0.5 μg for h*SOD1^WT^*/SOCK, 2.5 μg for *Ctr1^+/−^*/wild-type) were incubated with loading buffer (17.5% sodium dodecyl sulphate, 50% glycerol, 400 mM dithiothreitol (DTT), 0.3 M Tris-base (pH 6.8), 0.25% bromophenol blue) for 45 min at 56 °C to reduce and denature proteins, before being loaded onto 4–12% Bis–Tris Criterion pre-cast gels (Bio-Rad, Hercules, CA) and separated by sodium dodecyl sulphate–polyacrylamide gel electrophoresis in a Mini-PROTEAN Tetra Cell system at 180 V for 40 min at 4 °C (Bio-Rad). A loading control comprising equal amounts of protein from all four genotypes was also included in each gel. Separated proteins were transferred to Immobilon-PSQ PVDF (Millipore, Billerica, MA) membranes overnight at 9 V at 4 °C, before being air-dried and blocked in 5% skim milk (Bio-Rad, Hercules, CA) in phosphate buffer saline containing 0.1%Tween®20 (PBST) (Sigma-Aldrich, St. Louis, MO) for 1 h at room temperature. Membranes were then incubated overnight with primary antibodies against SOD1 (16 kDa, 1:2000) and GAPDH (35 kDa, 1:10,000) diluted in 1% skim milk in PBST overnight at 4 °C (**Supplementary Table 1**), followed by incubation with horseradish peroxidase (HRP)-conjugated goat anti-rabbit IgG (1:5000, Bio-Rad, Hercules, CA) for 2 h at room temperature. Protein signals were obtained using an ECL western blotting detection system (Bio-Rad, Hercules, CA) as per the manufacturer’s instructions, and developed using the iBright imaging system (Invitrogen). Signal intensities for SOD1 and GAPDH were quantified by densitometry using iBright analysis software v5.2.2 (Invitrogen), with SOD1 values normalized to corresponding GAPDH values for the same sample, as well as those for the loading control in each gel. GAPDH values were unchanged between genotypes at each age in all regions, validating the choice of this protein as a housekeeping gene (**Supplementary Figure 5**).

### Quantification of tissue metal levels

Metal levels in soluble and insoluble tissue extracts were quantified using our group’s established ICP-MS protocol^23^. Tissue homogenates (20-30 µL) were dried and digested overnight using concentrated nitric acid (50 µL, 70%, Suprapur grade, Merk Millipore) at room temperature, before being digested for a further 30 min at 70 °C, incubated for 1 h at 70 °C with concentrated hydrogen peroxide (30%, VWH International, PA, USA), and finally diluted to 2 mL with 1% nitric acid (1:10 v/v; Suprapur grade, Merk Millipore) prior to analysis. Total metal levels in each sample were measured in triplicate using a Perkin Elmer Nexion 300X Inductively Coupled Plasma Mass Spectrometer. Buffer controls containing 1% nitric acid were incorporated every 50 samples. Helium (4 mL/min) was used as a collision gas for the removal of polyatomic interferences. Measured mass-to-charge (*m/z*) ratios were 63 (Cu) and 66 (Zn). External calibration was performed using S24 multi-element standards (High Purity Standards, USA) diluted in 1% HNO_3_, while rhodium (Rh; m/z = 45) was used as a reference element via online introduction with a Teflon T-piece. Measurements were background-corrected to metal levels in buffer controls, adjusted for dilution factors and standardized against original wet tissue weights. All sample measurements were above the instrument’s limits of detection.

### SOD activity measurement

Total SOD activity was quantified in tissue extracts using a commercial SOD Assay Kit (Cat. #19160, Sigma-Aldrich, USA) according to the manufacturer’s instructions^9^. Briefly, samples containing 2 µg protein were diluted serially between 10- and 1000-fold and the assay signal measured in triplicate. A bovine SOD standard was used to generate a standard curve relating SOD activity to assay signal, which was applied to sample dilution curves to obtain SOD activity measurements. Total SOD activity in each sample was normalized to SOD1 protein levels measured using immunoblotting, which yielded a measure of SOD activity per unit of SOD1 protein in each sample.

### Fixed mouse tissue preparation for immunostaining

Mouse brain and lumbar spinal cord tissues for immunofluorescent staining were fixed overnight in 4% paraformaldehyde, before being incubated in 30% (w/v) aqueous sucrose solution in a 50 mL sample collection tube for 48 h. They were then embedded in Tissue-Tek® optimal cutting temperature medium (Sakura Finetek, Nagano, Japan; #4583), frozen at −20 °C and 30 µm thick serial tissue sections cut using an Epredia^TM^ CrytoStar^TM^ NX50 cryostat (ThermoFisher Scientific). These were divided into five free-floating section series and stored in anti-freeze solution (0.1 M PBS containing 30% glycerol and 30% ethylene glycol) at −30 °C until staining.

### Immunofluorescent staining and imaging

Free-floating brain and spinal cord tissue sections for immunofluorescent staining were brought to room temperature and washed with 0.3% Triton X-100 made in PBS (PBST) for 45 min to remove anti-freeze solution and increase tissue permeability, before antigen retrieval was performed using citrate buffer at 95 °C for 30 min (Vector Laboratories, USA). After blocking with 10% normal horse serum made in 0.1% PBST for 1 h at room temperature, sections were incubated with appropriate primary antibodies (**Supplementary Table 1**) at 4 °C for two nights. Following PBST washes, sections were incubated with appropriate non-spectrally overlapping fluorescent secondary antibodies (1:1000) and Hoechst nuclear staining dye (1 µg/mL) under the same conditions as the primary antibodies. Sections were then washed with PBST, mounted and coverslipped using Diamond Antifade Mountant (ThermoFisher Scientific, USA).

High throughput image acquisition was performed using a VS-200 Slide Scanner Microscope System (Olympus Life Science, Shinjuku-ku, Tokyo, Japan). Overview images of entire slides were first acquired at 10x magnification, utilizing ubiquitous Hoechst staining to correct for z-plane deviations. The SN (encompassing the SNc and SNr) was then manually outlined by G.S. for subsequent high-resolution scanning, which was performed by collecting 20 µm z-stacks at 63x magnification under oil immersion. Fluorescent signals were measured using a sequential acquisition paradigm, whereby longer wavelengths were excited and captured first; 647 nm, 568 nm, 488 nm, 405 nm. A no primary negative control was run to validate the absence of non-specific fluorescent signals, whereby primary antibodies were substituted for normal serum, which produced no detectable fluorescent signal using the same acquisition settings as experimental samples (**Supplementary Figure 6**).

### Immunofluorescent image analysis

We developed a novel image analysis workflow to count dopamine neurons and measure the volume of disSOD1 and astrocytes in the SNc of all mouse strains. Image analyses were performed in a manner that was blinded to mouse genotype and treatment, with images assigned a random number by a secondary investigator (V.C.). This workflow consisted of three stages: pre-analysis image preparation, retraining an existing artificial intelligence (AI) deep learning model (Cellpose) to segment dopamine neurons, and image analysis (**Supplementary Figure 7**). All ImageJ and Python scripts used to collect data are publicly available on GitHub (owner: Richard Harwood, repository: <u>Sod1_CuATSM_Image_Analysis</u>).

#### Pre-analysis image preparation

The pre-analysis stage functioned to 1) crop z-stack images to only contain the SNc and SNr, 2) generate masks outlining the SNc and SNc+SNr in each cropped z-stack, and 3) identify suitable intensity thresholds for SOD1 immunostaining in each z-stack to enable delineation of SOD1 protein aggregates from background tissue fluorescence. Briefly, raw z-stack images generated by the VS200 Slide Scanner Microscope (.vsi files) of tissues multi-stained for disSOD1, TH, GFAP and Hoechst were cropped by G.S. broadly into boxes containing the entirety of the SN using the open-source software platform, QuPath (v.0.5.1). Resulting .ome files of these regions (no compression) were then processed in ImageJ using a custom macro script (*prepare_masks_and_other_processing.ijm*), which converted them to .tiff files and 1) set the brightness and contrast of all images to a defined grayscale intensity (115-700), 2) prompted the investigator (G.S.) to outline two specific SN regions of interest within the broader SN - SNc, SNc+SNr - before converting them to a binary-filled mask, and 3) prompted the investigator (G.S.) to manually set disSOD1 signal thresholds for each image to account for variable stain intensity between runs (**Supplementary Figure 7**). Ten percent of these images were re-thresholded and masks re-applied by the same investigator (G.S.) to evaluate the intrarater reliability of data obtained from these images (Cronbach’s alpha = 0.986, n = 8 mice), before being subject to independent segmentation and counting by another investigator (V.C.) to measure the interrater reliability of these data (Cronbach’s alpha = 0.988, n = 8 mice).

#### Retraining AI deep learning model for neuron segmentation

Accompanying the processing of raw z-stack images, we trained an open-source deep learning AI Cellpose model (*sod1_cellpose.ipynb*) to segment and quantify dopamine neurons. Training was performed using 50 individual 2D slices from 16 z-stack images (8 different mice; 10% of cohort), which had been manually segmented and counted by one investigator (G.S.). Twelve percent of all images were re-segmented and recounted by the same investigator (G.S.) to evaluate the intrarater reliability of data used to train the Cellpose model (Cronbach’s alpha = 0.997, n = 6 mice), before being subject to independent segmentation and counting by another investigator (V.C.) to measure the interrater reliability of these data (Cronbach’s alpha = 0.978, n = 6 mice). The model was trained using 500 epochs with a learning rate of 0.2 using the “Cytoplasm2” as the starting weights. Whilst the model was trained on 2D images 3D cell segmentation was achieved by merging segmented immunopositive staining throughout the z-plane, provided that the intersect over union (IoU) between the mask on the current slice and the next slice is greater than or equal to 0.25. The resulting 3D images were then binarized and cleaned using dilation, fill holes, erosion and remove small objects functions and relabeled as discrete cells. Following training, we tested the accuracy of the model by applying it to an additional set of eight 2D z-slices from z-stacks in which neurons had already been manually segmented. A common way to assess the difference between ground truth data and predicted data is an F1 score which is calculated from a confusion matrix, which captures the True/False positives and negatives. The F1 score is calculated as:

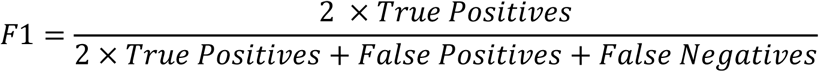

Our model produced an F1 score of 0.83. We also compared individual cell counts from the ground truth and predicted data observing a strong correlation (R^2^ = 0.9197; **Supplementary Figure 8).** These results are comparable in performance to previous studies utilizing AI tissue-based cell segmentation for images with an increased signal-to-noise ratio^24, 25^.

#### Image analysis

The analysis stage employed a custom Python script (*CuATSM_treatment_sod_1_loop.ipynb*), which was run through Jupyter software, to process the .tiff images generated during pre-analysis image preparation. This workflow generated a spatial profile of SOD1, TH and GFAP immunostaining within individual z-slices of a given z-stack, which were then combined to form a 3D reconstruction incorporating all three staining types and their relative positions in space. These representations were restricted to regions of interest (SNc only, SNc+SNr) by applying the pre-defined masks and SOD1 intensity thresholds defined earlier, with quantification of the volume of each of these components also made possible by counting pixel numbers in each slice and converting them to voxels upon 3D reconstruction (1 voxel = 0.125 µm^3^). This allowed estimation of the volume of SOD1 aggregates within and outside of TH- or GFAP-positive immunostaining to estimate the cellular localization of this pathology.

Neuron stereology was performed alongside volumetric analyses by our custom Python script, which applied SNc masks to .tiff z-stack images and ran them through the retrained AI Cellpose model to obtain the neuronal count. These were converted to neuron densities by accounting for the volume of the SNc in which they were counted.

The complex and heterogeneous morphology of astrocytes, together with the frequent overlap of their projections, prevented accurate astrocyte counts from being generated in this study. To capture the non-uniform morphology of an astrocyte for volumetric analyses, our custom Python workflow applied triangular yen thresholding, adding Gaussian blur and size filters to remove small objects. Twenty-five grayscales were also removed from the background signal to separate cells from background and enable estimation of astrocyte volume in the SNc and SNr.

Finally, each z-stack analysed with our custom Python code generated a separate csv file, which we combined using a second Python script (*sod_1_csvs.ipynb*) for subsequent statistical analyses.

### Mouse lumbar spinal cord motor neuron stereology

Spinal motor neuron stereology was performed in mouse lumbar spinal cord sections (50 sections/mouse) using immunofluorescent staining for the motor neuron markers choline acetyltransferase (ChAT) and Islet-1 (ISL-1)(**Supplementary Table 1**)(**Supplementary Figure 9**) as described above, with imaging conducted on an Olympus VS200 virtual slide scanner (Olympus, Japan) at 40x magnification using the maximum intensity projection function. Spinal cord sections were confirmed as being from the lumbar region using anatomical features identified in an atlas of the mouse spinal cord^26^. Once images were obtained, left and right grey matter horns that were ventral to the spinal canal were then converted to .tiff files separately using QuPath (v0.5.1) software. Ventral motor neurons that were considered positive for both ISl-1 and ChAT were then segmented for counting using a custom ImageJ script, as previously described^13^. This script is publicly available on GitHub (owner: Richard Harwood, repository: <u>Sod1_CuATSM_Image_Analysis</u>). Motor neurons were defined as cells that possessed nuclear ISL-1 staining with proximal ChAT staining above background signal intensity. Counts were normalized to a 1 mm length of spinal cord by dividing the counts by the combined length of spinal cord comprised by the sections counted (e.g. 50 sections of 30µm thickness = 1.5 mm length of spinal cord). Manual segmentation of ventral spinal cord motor neurons was performed on 10% of the dataset to confirm the reliability of automated data produced, whereby motor neurons that were positive for both ISL-1 and ChAT staining were counted in the extracted images. Manual counting demonstrated excellent reliability for the automated segmentation (Cronbach’s α= 0.979 n= 70).

### Quantification of striatal dopamine levels and turnover

Fresh-frozen striatal mouse tissue was homogenized by pulse sonication (40% duty cycle, output 4, 2 × 30 sec intervals) in a solution of 150mM glacial acetic acid and 500 μM diethylene triamine penta-acetic acid dissolved in LC-MS grade water (19x vol (uL) per tissue weight (mg)), which was spiked with 1 μM Salsolinol hydrobromide as an internal standard. Homogenized tissue was then centrifuged at 14,000 *g* for 25 min at 4 °C and the supernatant collected. Ten microlitres of supernatant was subjected to protein determination using a bicinchronic acid assay (ThermoFisher, USA), while the remaining supernatant was spun in 3 kDa cut-off Amicon® centrifugal filter tubes (Merck Millipore) at 14,000 *g* for 90 min at 4 °C and filtrates collected and stored at −80 °C for analysis.

Quantification of dopamine, 3,4-dihydroxyphenylacetic acid (DOPAC), and homovanillic acid (HVA) levels were conducted using a reverse-phase high performance liquid chromatography (HPLC) system, consisting of a pump module (Shimadzu Prominence LC-20AD, Shimadzu Corporation) coupled to a reversed phase Gemini C18 110 Å column (5 μm pore size, 150 × 4.6 mm; Phenomenex) and an electrochemical detector (Antec Leyden) with a glassy carbon-working electrode maintained at +0.802 V against a Ag/AgCl reference electrode. Both detector and column were maintained at 40 °C. Twenty microlitres of prepared samples were injected via a Prominence autosampler (Shimadzu Prominence SIL-20A, Shimadzu Corporation, Kyoto, Japan). The mobile phase comprised of 13% (v/v) methanol, 8.7 mM sodium phosphate monobasic, 100 μM ethylenediaminetetraacetic acid, 0.6 mM octane sulfonic acid and 0.05 mM triethyl amine in MilliQ water, maintained at a pH of 2.81 and an isocratic flow. Samples were injected at a volume of 15 µL, with a flow rate of 1.5 mL/min and a 15 min total run time. Calibration curves prepared from starting concentrations of analytical standards of dopamine (1 μM), DOPAC (0.5 μM), HVA (2 μM) and Salsolinol (1 μM) in homogenization buffer were run at the beginning of each day to quantify any variability in the HPLC system. The Shimadzu integrated workstation LabSolutions software (Version 5.57; Shimadzu) was used to calculate the area under the curve for each peak of interest, and data subsequently normalized to 1) the total protein concentration of each sample, and 2) the concentration of internal standard Salsolinol. Dopamine turnover was calculated as the ratio of the sum of HVA and DOPAC concentrations to that of dopamine. Neurotransmitter concentrations are expressed as ng/mg of protein (mean ± SEM).

### Animal locomotor function

Animals were subjected to a battery of motor performance tests at 5.5 months-of-age following the completion of vehicle/CuATSM treatments (**Supplementary Figure 10)**.

#### Inclined balance beam

Motor coordination and balance were assessed using an inclined balance beam test as previously described^27^. Mice were trained over two days using four to five practice trials per day, where they were required to traverse increasing lengths of an elevated round beam (14 mm in diameter, 60 cm total length) to reach the shelter of an enclosed goal box. Mice were acclimated to the testing environment over two days, spending 30 minutes in the room and an additional 2 minutes in the goal box each day prior to training. To prevent premature exploration, the goal box opening was temporarily covered if a mouse attempted to exit. During training, mice were initially placed on the balance beam just outside the goal box, with the starting distance progressively increased in 10 cm increments until they could traverse the full 60 cm unassisted. Each training run was followed by a 1-min rest period in the goal box. On the third day, data collection commenced with four full-length trials, each separated by a 30-second rest interval. Endpoints were either reaching the goal box or falling from the beam (height 1m). All trials were video recorded using two cameras placed at 45° either side of the balance beam for subsequent analysis. Measurements recorded for analysis included the time taken to traverse the beam (latency) and the number of times the feet slipped from the beam while traversing (paw slips). Mice that fell immediately after being placed on the beam were returned to the same position for re-testing, whereas those that fell after walking a short distance up the beam were returned to the goal box to rest for 30 sec. Recordings of incomplete beam crossings were not analyzed, with all animals tested until they had completed four full beam crossings.

#### Grip strength

Mouse forelimb grip strength was measured using a Digital Force Gauge (Chatillon, USA). Mice were positioned close to a horizontal bar that was attached to the force gauge, allowing them to reach and grab onto the bar with their forelimbs. Mice were then positioned so that their body was horizontal and in line with the bar, and were pulled horizontally away from the bar by the tail until they released their grip. The tension was measured and defined as grip strength, which was normalized to mouse body weight. Mice underwent testing 4 times and were rested in their cages with food and water for 1 min between trials.

#### Open field

Spontaneous motor activity of mice was assessed using an open field test, in which mouse locomotion within a 400 mm x 400 mm box was recorded using two mounted cameras. Mice were allowed to freely explore for 10 min, after which time they were returned to a new cage, as returning them to their home cage directly after testing can affect exploratory behaviour. Measures of interest included total distance travelled, average speed, time spent immobile, immobile episodes, entries into central zone (20 mm x 20 mm), which were all obtained by processing video footage using ANY-maze Video Tracking System software (version 7.2; ANY-maze, Wood Dale, IL, USA).

### Statistical analyses

Statistical analyses were performed using SPSS 28.0 (IBM Corp, NY, USA). Outliers were defined as ≥3× the interquartile range and excluded from the analysis. Parametric tests or descriptive statistics with parametric assumptions (standard one- and two-way ANOVA) were used for variables meeting the associated assumptions, with data normality assessed using the Shapiro-Wilk test, and homogeneity of variance was evaluated using Levene’s and Brown– Forsythe tests. One-way and two-way analysis of variance (ANOVA) were paired with Dunnett’s multiple comparisons post hoc test to assess pairwise comparisons between experimental and control groups for a given variable. Non-parametric statistics (Spearman’s correlation) were used for variables where the observed data did not fit the assumptions of parametric tests. Where large differences between groups existed for biological reasons (e.g. SOD1 protein expression, SOD1 proteinopathy), data were log transformed prior to the application of statistical tests. Cronbach’s alpha was used to measure interrater reliability between researchers performing quantitative stereology. Significance level was defined as p < 0.05 for all statistical tests. Graphs were generated using GraphPad Prism 9.4.0 (Graph-Pad, CA, USA).

## Results

### CuATSM mitigates the development of Parkinson-like disSOD1 pathology

Oral administration of 30 mg/kg CuATSM daily for 4-to-6 months increases spinal cord copper content in transgenic mutant SOD1 mice^14, 15, 28^. Here, we report oral administration of 15 mg/kg CuATSM for as little as 3 weeks increased brain copper 2-fold higher than physiological levels, while treatment with 30mg/kg CuATSM resulted in a 3-fold increase in brain copper, raising concerns about potential toxicity (**Supplementary Figure 11**). To reduce risk of copper toxicity from chronic CuATSM treatment^29^, we therefore treated SOCK mice with either 15 mg/kg CuATSM or vehicle (*n* = 10/treatment) each day from weaning until 5.5 months-of-age to examine whether CuATSM could mitigate age-associated development of Parkinson-like wild-type SOD1 pathology, which is present by 6 weeks of age.^13^ Wild-type mice, as well as both mouse strains crossed to generate SOCK mice (copper deficient *Ctr1^+/−^* mice, human wild-type SOD1 overexpressing h*SOD1^WT^* mice), were treated in parallel as controls (*n* = 10/genotype/treatment). The volume of SOD1 pathology in the SN of these animals was then quantified using 3D reconstructions of SOD1 immunostaining, which utilized a conformation-specific SOD1 antibody selectively recognizing structurally-disordered (dis)SOD1. These analyses encompassed the dopamine neuron-rich SN pars compacta (SNc; **Fig. 1a**), as well as the astrocyte-rich SN pars reticulata (SNr; **Fig. 1b**), with disSOD1 pathology separately quantified within, and outside, of dopamine neurons and astrocytes in these regions (**Fig. 1c**). Consistent with our published description of the development of disSOD1 pathology in these four mouse strains^13^, disSOD1 accumulation was significantly greater within the SN of vehicle-treated SOCK mice, compared with vehicle-treated h*SOD1^WT^* (3.6-fold increase), *Ctr1^+/−^* (7.2-fold increase) and wild-type (19.7-fold increase) mice (**Fig. 1d**). Remarkably, CuATSM treatment elicited a 4.4-fold decrease in the volume of disSOD1 deposits in the SN of SOCK mice (**Fig. 1d**), reducing the volume of this pathology below that in the SN of vehicle-treated h*SOD1^WT^*control mice. In all mouse strains, the concentration of disSOD1 in the SN was inversely correlated with midbrain copper levels (**Fig. 1e,f**), but not tissue zinc levels, which were unchanged between vehicle and CuATSM treated animals for each strain (**Supplementary Figure 12**). These data support a role for copper in the development of wild-type disSOD1 pathology in SOCK mice. Interestingly, the volume of disSOD1 was not significantly reduced following CuATSM treatment in h*SOD1^WT^* mice (**Fig. 1d**), suggesting the lower quantities of disSOD1 pathology observed in h*SOD1^WT^* mice, as well as residual pathology in CuATSM-treated SOCK mice, may be driven in part by copper-independent mechanisms.

**Figure 1.**
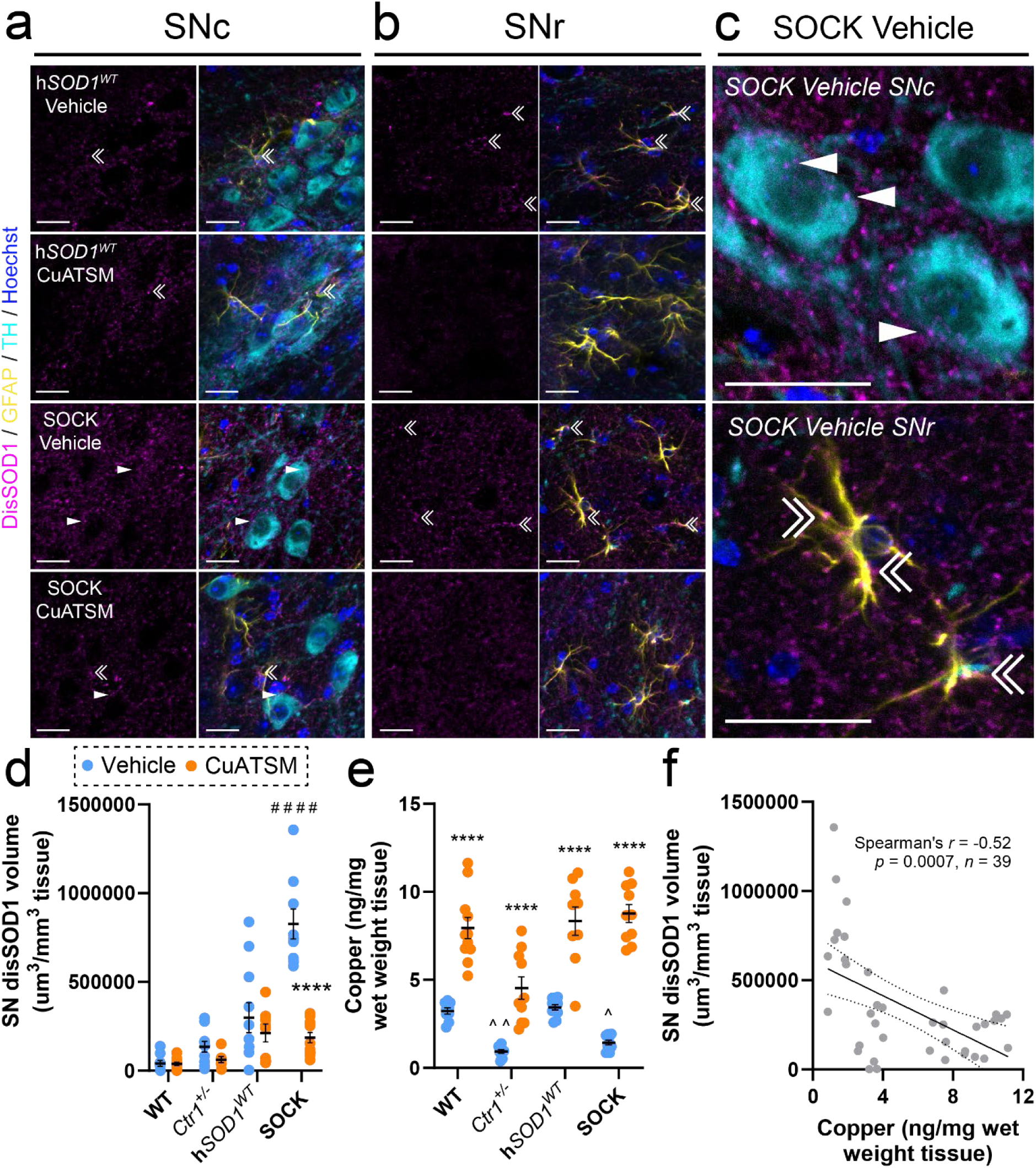
Quantification of total disSOD1 burden in the SN of all mouse strains following treatment. Immunofluorescent staining of fixed midbrain tissues from the substantia nigra pars compacta (SNc) and substantia nigra pars reticulata (SNr) of h*SOD1^WT^*and SOCK mice treated daily with vehicle or 15 mg/kg CuATSM. Immunostaining utilized the unfolded beta barrel (UβB) conformation-specific SOD1, which revealed disSOD1 aggregates (magenta) within and outside of dopamine neurons (TH, cyan; white arrowheads) and astrocytes (GFAP, yellow; double white arrowheads). Individual panels for immunostaining of the SNc and SNr of SOCK and h*SOD1^WT^* mice, as well as *Ctr1^+/−^* and wild-type mice, are presented in **Supplementary Figures 13 and 14**. Antibody details are presented in **Supplementary Table 1**. Scale bars represent 20 µm in panels **a-c. d.** The volume of disSOD1 expressed relative to the total volume of tissue within which it was quantified varied significantly between genotypes and was elevated in SOCK mice compared with h*SOD1^WT^* mice treated with vehicle. CuATSM treatment elicited a significant decrease in this pathology in SOCK mice but not in any control mouse strains. **e.** Midbrain copper levels were decreased in vehicle-treated *Ctr1^+/−^* and SOCK mice compared with wild-type mice, while CuATSM treatment induced significant increases in midbrain copper levels in all four mouse strains. Data in panels **d** and **e** represent mean ± SEM (*n* = 10/genotype/treatment), with full details of statistical tests presented in **Supplementary Table 2**. Comparisons marked with an asterisk (*) denote those made between vehicle- and CuATSM-treated mice of the same genotype, those marked with an arrowhead (^) demarcate those made to vehicle-treated wild-type mice, while those marked with a hashtag (#) denote those made to vehicle-treated h*SOD1^WT^* mice. #### *p* < 0.0001, \*\*\*\**p* < 0.0001, ^*p* < 0.05, ^^*p* < 0.01. **f.** DisSOD1 volume in the SN was inversely correlated with midbrain copper content in vehicle- and CuATSM-treated SOCK and h*SOD1^WT^* mice. Statistical test details are presented in the panel.

### Astrocytes accumulate disSOD1 pathology and are disproportionally impacted by CuATSM treatment

Previous characterization of disSOD1 pathology in the SOCK mouse SN revealed minimal (∼2%) colocalization with dopamine neuron soma^13^, implicating additional cell types in the development of this pathology. Among these, astrocytes emerge as a key potential mediator, given they play essential roles in brain metal storage and homeostasis^30^, and exhibit SOD1 inclusions in transgenic mutant SOD1 mice^31^ and *SOD1*-linked ALS patient tissues^32^. To evaluate the relative involvement of astrocytes and neurons in the development of disSOD1 pathology, we incorporated immunostaining for glial fibrillary acidic protein (GFAP, astrocytes) and tyrosine hydroxylase (TH, dopamine neuron soma) into 3D reconstructions of disSOD1 pathology in the SN of all four mouse strains (**Fig. 2a, Supplementary Figure 15**), and quantified the proportion of pathology within, and outside, of both cell types in vehicle-treated animals (**Fig. 2b, Supplementary Table 3**). Astrocytes contained 28.2% of the total volume of disSOD1 pathology across the SN of SOCK mice, 75% of which was localized to the glia-rich SNr (**Fig. 2b**). Importantly, astrocytic disSOD1 comprised a lower proportion (15.8%) of the total volume of this pathology in the SN of vehicle-treated h*SOD1^WT^* mice (**Fig. 2b**), suggesting a proportionally greater accumulation of disSOD1 in astrocytes in SOCK mice. Indeed, astrocytic disSOD1 volume was increased 4.9-fold in vehicle-treated SOCK mice compared with h*SOD1^WT^*mice, which is significantly higher than the 3.5-fold increase in this pathology outside of dopamine neuron soma and astrocytes between these strains (**Fig. 2b**). Increases in astrocytic disSOD1 were associated with a greater volume of astrocytes across the SN of vehicle-treated SOCK mice compared with h*SOD1^WT^* and wild-type mice (**Supplementary Figure 16**), which is consistent with disSOD1-induced astrogliosis observed in transgenic mutant SOD1 mice^33^. Collectively, these data identify astrocytes as a key site for the development of disSOD1 pathology in the SOCK mouse SN.

**Figure 2.**
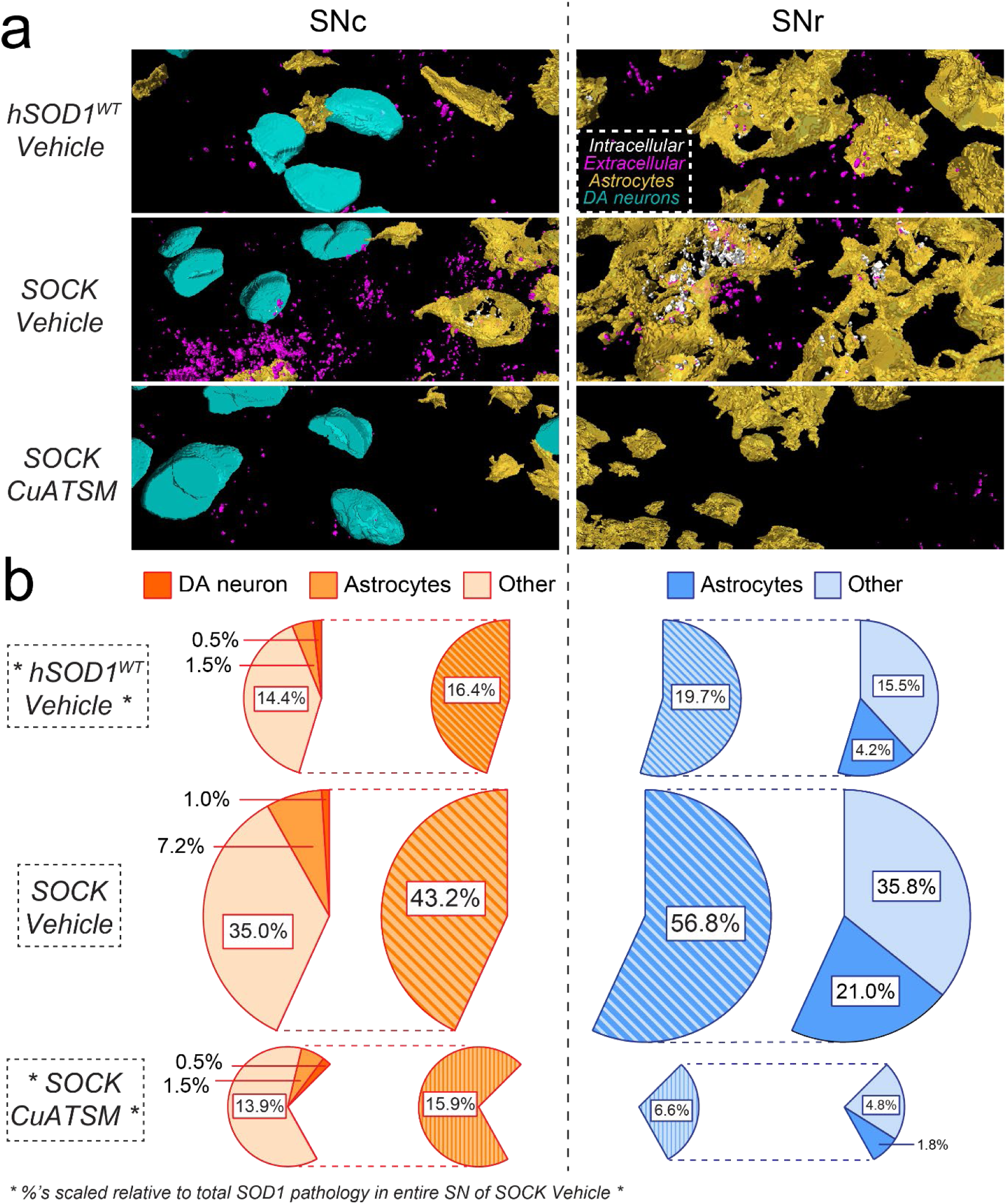
Impact of CuATSM treatment on the distribution of disSOD1 pathology in the SN of SOCK mice. **a.** Three-dimensional reconstructions of immunostaining for disSOD1 (UβB), dopamine neurons (TH, cyan) and astrocytes (GFAP, yellow) in the SNc and SNr of SOCK and h*SOD1^WT^* mice. DisSOD1 localized within or outside of either cell type is presented in white or magenta, respectively. Scale bars represent 20 µm. Reconstructions of all four mouse strains treated with vehicle or CuATSM are presented in (**Supplementary Figure 15**). Panels displayed in this figure constitute those most important for illustrating the higher level of disSOD1 pathology in SOCK mice and the impact of CuATSM treatment. **b.** Proportions of disSOD1 pathology colocalized within dopamine (DA) neurons, astrocytes, or other compartments (other). Percentages were generated by scaling the raw volume of disSOD1 in each compartment to the total volume of disSOD1 in the entire SN (SNc + SNr) of vehicle-treated SOCK mice, as this represents the maximum disSOD1 volume exhibited by any mouse strain/treatment combination. Raw disSOD1 volumes are reported in **Supplementary Table 3**.

Astrocytic disSOD1 pathology was significantly attenuated following CuATSM treatment; astrocytic disSOD1 volume in the SN decreased 8.5-fold in CuATSM-treated SOCK mice compared with that in vehicle-treated SOCK animals (**Fig. 2b**). This effect was far greater than the 2-fold decrease in disSOD1 volume observed within dopamine neuron soma, as well as the 3.8-fold decrease in this pathology outside of dopamine neurons and astrocytes, which may explain why CuATSM mitigated disSOD1 development in the glia-rich SNr to a greater extent (8.6-fold decrease) than the neuron-rich SNc (2.7-fold decrease) (**Fig. 2b**). These alterations were not associated with changes to astrocyte volume in SOCK mice, contrast to the significant increase in SN astrocyte volume in wild-type and h*SOD1^WT^* mice following CuATSM treatment (**Supplementary Figure 16**). Taken together, our data suggests that CuATSM mitigates the development of Parkinson-like disSOD1 pathology in the SN of SOCK mice, particularly within astrocytes.

### CuATSM treatment largely restores SOD1 post-translational modifications in SOCK mice

The development of disSOD1 pathology is associated with alterations to SOD1 post-translational modifications (PTMs)^13^. These include higher levels of atypical modifications that promote SOD1 misfolding and dysregulation of physiological modifications that normally regulate the structure, maturation, localization and function of SOD1. To determine whether CuATSM mitigates these atypical alterations, we employed proteomic mass spectrometry to profile the post-translational fingerprint of SOD1 protein immunoprecipitated from the SN of vehicle- and CuATSM-treated SOCK mice. These were compared with SOD1 PTMs in the SN of vehicle-treated h*SOD1^WT^* mice to ensure alterations did not simply derive from increased SOD1 protein expression. Consistent with our previous study, we identified 40 altered SOD1 PTMs across 33 residues (21.4%) of SOD1 protein in vehicle-treated SOCK mice compared with h*SOD1^WT^*mice, including increases in oxidation (**Fig. 3a)** and glycation, as well as dysregulation of SOD1 glycosylation (**Fig. 3b,c)**, acetylation, succinylation, deamidation, ubiquitylation and phosphorylation (**Fig. 3d**, **Supplementary Table 4**)(**Table 1**). Among these was significant oxidation of all four copper-binding histidine residues (**Fig. 3a)**, as well as glycation of a solvent-accessible arginine and lysine residue, all of which underlie aggregation of the protein *in vitro* by promoting SOD1 demetallation and misfolding^34, 35^. This is likely to be compounded by dysregulation of SOD1 protein trafficking following decreased glycosylation (**Fig. 3b,c)**^36^, as well as disruptions to its turnover from decreased deamidation and ubiquitylation^37, 38^, with lower levels of acetylation, succinylation, phosphorylation further impacting the maturation and catalytic activity of the protein^37, 38^. Astonishingly, CuATSM treatment restored 29 (73%) of the 40 PTM alterations in SOCK mice to levels observed in h*SOD1^WT^* control mice (**Fig. 3e**, **Supplementary Table 4**) (**Table 1**). These included atypical and physiological PTMs, with decreases in histidine oxidation and restoration of SOD1 glycosylation exhibiting the greatest magnitudes of change. The levels of a further 4 (10%) PTM alterations (oxidation H63, glycation K30, phosphorylation S25, deamidation N139) were also positively impacted by CuATSM treatment, shifting 2-to-8-fold closer to baseline levels observed in h*SOD1^WT^* mice compared with vehicle-treated SOCK mice (**Fig. 3e**, **Supplementary Table 4**). Given the clear role of these PTM alterations in promoting the development of disSOD1 pathology, we speculate that their correction underlies the decrease in disSOD1 volume in the SOCK mouse SN following CuATSM treatment.

**Figure 3.**
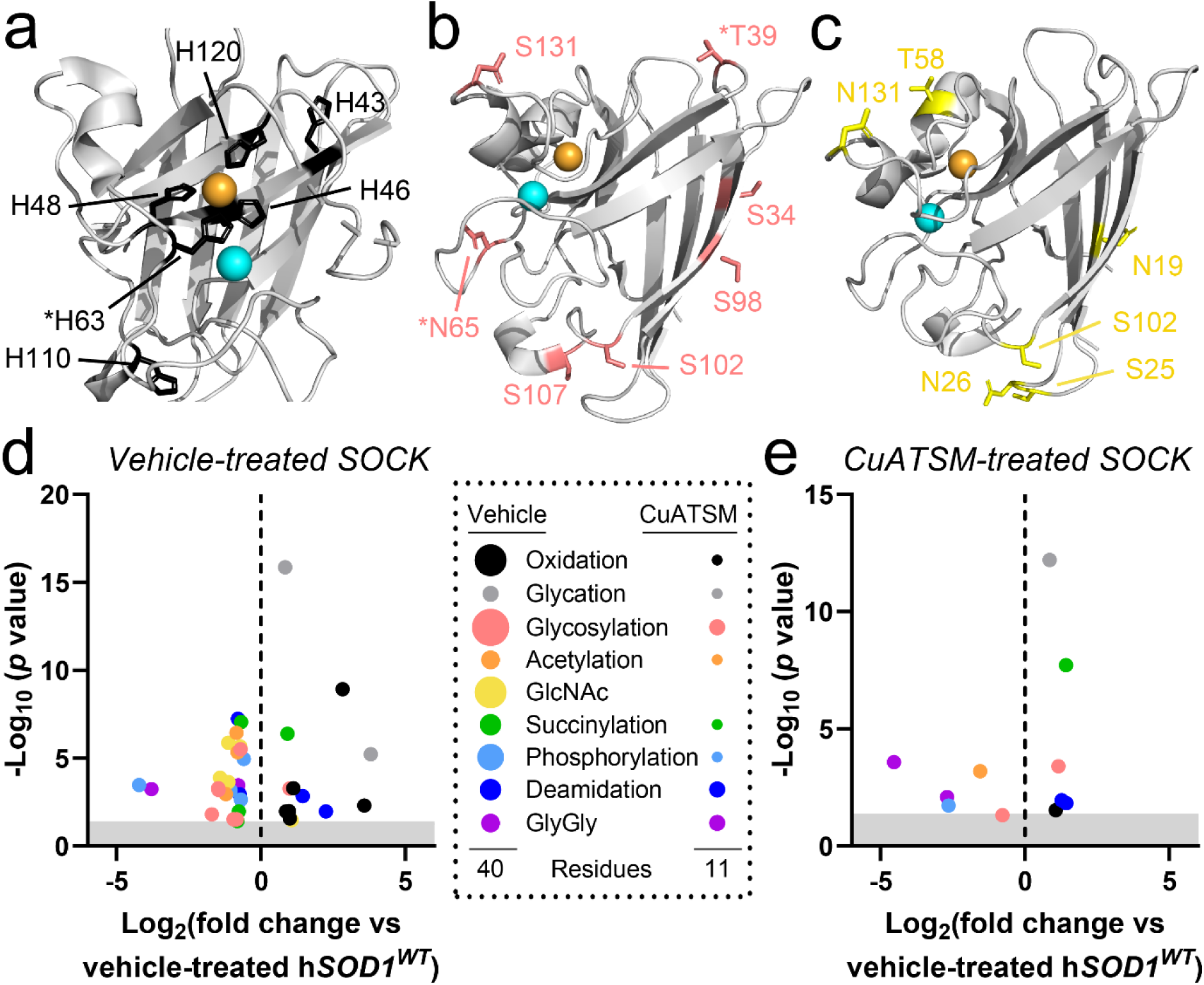
Altered SOD1 PTMs in the SN of vehicle- and CuATSM-treated SOCK mice. Atypical oxidation of SOD1 histidine residues was increased in the SN of SOCK mice compared with h*SOD1^WT^* mice (**a**), while glycosylation (**b**) and acetylglucosamine (**c**) were significantly decreased. Side chains of labelled residues are highlighted in black, red and yellow, respectively. Residues are labelled using one letter amino acid codes. Copper and zinc ions are highlighted in orange and cyan respectively. Alterations to SOD1 PTMs in vehicle-treated (**d**) and CuATSM-treated (**e**) SOCK mice. Circles for each type of modification in the legend are scaled to the number of residues where that PTM is altered in each treatment group. GlyGly modifications result from tryptic digestion of ubiquitin-conjugated proteins, which serve as indicators of protein ubiquitination. Complete details of statistical analyses identifying PTM alterations in vehicle- and CuATSM-treated SOCK mice are presented in **Supplementary Table 4**. Abbreviations: GlcNAc, acetylglucosamination.

**Table 1.** Known effects of altered SOD1 PTMs on the structure and function of the protein. Specific residues and magnitudes of alterations are outlined in Supplementary Table 4.

| Class | Modification | Known effects on SOD1 protein | Alteration in vehicle-treated SOCK vs vehicle-treated *hSOD1<sup>WT</sup>* mice | Alteration in CuATSM-treated SOCK vs vehicle-treated *hSOD1<sup>WT</sup>* mice |
| --- | --- | --- | --- | --- |
| **Increased atypical modification** | Oxidation | Dissociation of SOD1-bound Cu and Zn<sup>39</sup><br>Covalent crosslinking and aggregation of wild-type SOD1<sup>40, 41</sup> | 6 residues increased | 1 residue increased |
|  | Glycation | Protein unfolding and loss of secondary structure<sup>42</sup><br>Altered superoxide guidance and perturbed SOD1 activity<sup>43</sup> | 2 residues increased | 1 residue increased |
| **Dysregulated physiological modification** | Acetylation | Inactivates SOD1 by disrupting binding to CCS<sup>44</sup><br>Reduces SOD1 protein net charge (+1 to 0) and impedes electrostatic guidance of superoxide<sup>45</sup> | 3 residues decreased | 1 residue decreased |
|  | Succinylation | Reduces SOD1 protein net charge (0 to -1) and impedes electrostatic guidance of superoxide<sup>46</sup> | 3 residues decreased<br>1 residue increased | 1 residue increased |
|  | Phosphorylation | Impedes SOD1-mediated transcription of protective antioxidant genes by impairing SOD1 nuclear translocation<sup>47</sup> | 4 residues decreased<br>1 residue increased | 1 residue decreased |
|  | Deamidation | Impaired coupling of CCS–SOD1 and thermodynamic destabilization of SOD1 protein structure<sup>48</sup> | 2 residues decreased<br>2 residues increased | 2 residues increased |
|  | Ubiquitylation | Attempted degradation of aggregated SOD1 by the ubiquitin-proteasome system<sup>49, 50</sup> | 3 residues decreased | 2 residues decreased |
|  | Glycosylation<br>Acetylglucosamine | Associated with degeneration of spinal cord motor neurons in mutant *SOD1<sup>G93A</sup>* mice<sup>51</sup> | 11 residues decreased<br>2 residues increased | 1 residue decreased<br>1 residue increased |

**Abbreviations:** AGE, advanced glycation end-product; SOD1, superoxide dismutase 1; CCS, copper chaperone for SOD1; SOCK, SOD1 overexpressing *Ctr1* knockdown; PD, Parkinson disease; Ct, control, SN; substantia nigra.

### Improved antioxidant function of SOD1 protein following CuATSM treatment

None of SOD1’s copper-binding histidine residues are solvent-accessible when copper is bound, hence increased oxidation of these residues indicates decreased copper binding in SOCK mice compared with h*SOD1^WT^* mice^37, 38^. As copper binding is critical for the catalytic role of SOD1, the observed decrease in total SOD activity (**Fig. 4a**)^13^, as well as SOD activity per unit of SOD1 protein (**Fig. 4b**)^13^, in SOCK mice compared with h*SOD1^WT^* mice also suggests decreased levels of copper binding in these animals. The decrease in oxidation of SOD1’s copper-binding histidine residues in SOCK mice following CuATSM treatment (**Fig. 3e**, **Supplementary Table 4**) therefore suggests SOD1 copper binding is increased by this treatment, consistent with data from transgenic mutant SOD1 mice treated with CuATSM^14, 15^. Aligning with these data, CuATSM treatment also elicited higher total SOD activity (**Fig. 4a**) and restored SOD activity per unit of SOD1 protein in transgenic SOCK and h*SOD1^WT^* mice to levels observed in wild-type mice (**Fig. 4b**). This was achieved without alterations to SOD1 protein levels, contrasting non-transgenic wild-type and *Ctr1^+/−^* mouse strains, which exhibited an up-regulation of SOD1 protein expression alongside increases in total SOD activity (**Fig. 4e,f**). Furthermore, increases in SOD activity per unit of SOD1 protein were strongly correlated with tissue copper content in transgenic strains (**Fig. 4c**), suggestive of a rectification of the imbalance between the demand for, and supply of, copper to SOD1 in these mice. This is reinforced by the significant negative correlation between SOD activity per unit of SOD1 protein and concentration of disSOD1 in the SN of SOCK mice (**Fig. 4d**), both of which are heavily dependent on adequate SOD1 copper loading. Collectively, our data provides the first empirical evidence that the correction of altered SOD1 PTMs *in vivo* may mitigate the development of wild-type SOD1 pathology.

**Figure 4.**
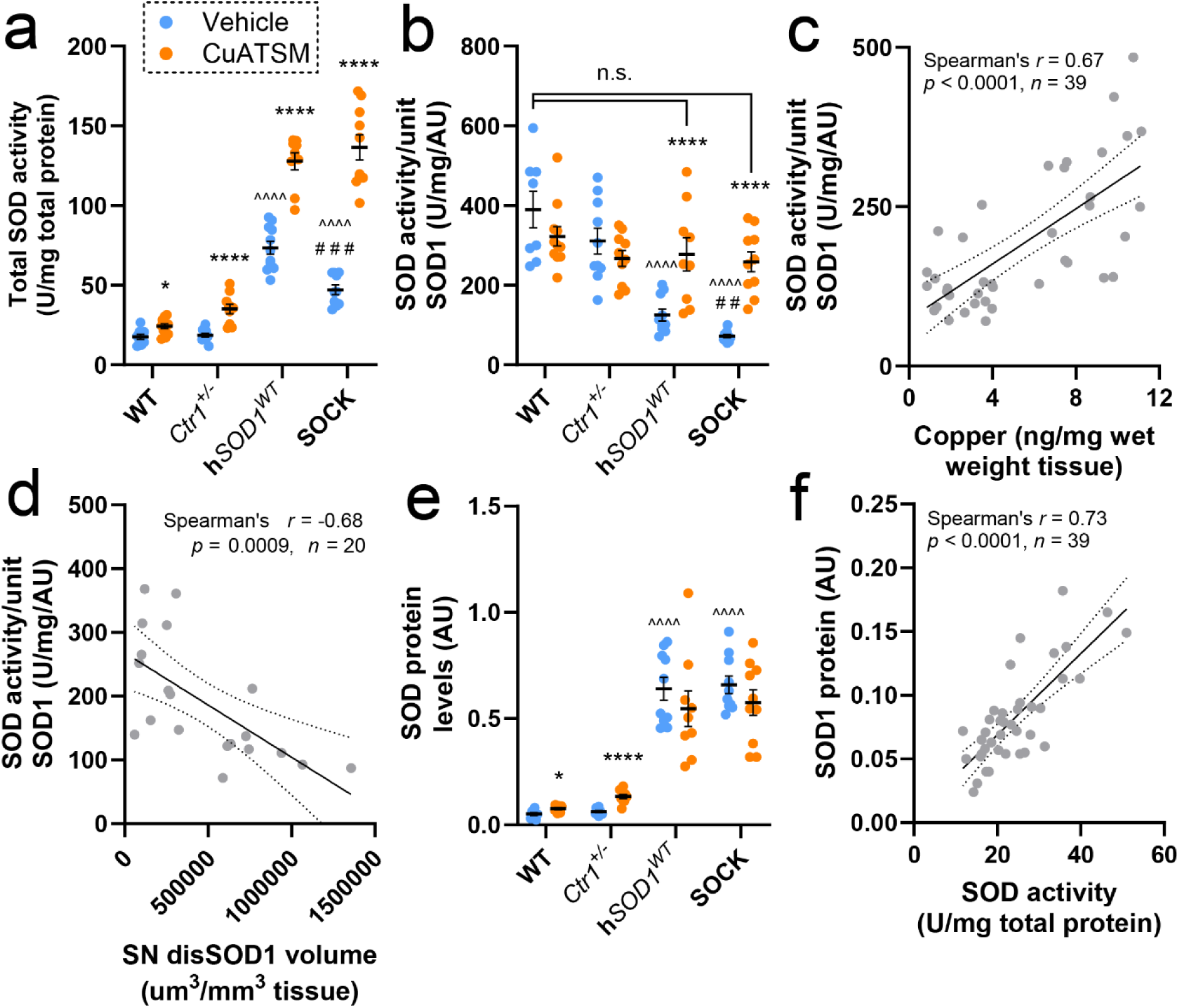
SOD1 enzymatic activity and protein expression in the SN of all mouse strains following treatment. **a.** Total SOD activity varied significantly between mouse strains and was increased in vehicle-treated SOCK and h*SOD1^WT^*mice compared with wild-type (WT) and *Ctr1^+/−^* mice, as well as vehicle-treated SOCK mice compared with h*SOD1^WT^* mice. CuATSM treatment elicited increases in total SOD activity across all strains. **b.** SOD activity per unit of SOD1 protein varied significantly between mouse strains and was decreased in vehicle-treated SOCK and h*SOD1^WT^* mice compared with wild-type mice, as well as vehicle-treated SOCK mice compared with h*SOD1^WT^* mice. CuATSM treatment increased SOD activity per unit of SOD1 protein in SOCK and h*SOD1^WT^*mice. **c.** SOD activity per unit of SOD1 protein was positively correlated with midbrain copper content in vehicle- and CuATSM-treated SOCK and h*SOD1^WT^* mice. **d.** DisSOD1 volume in the SN was inversely correlated with SOD activity per unit of SOD1 protein in vehicle- and CuATSM-treated SOCK mice. **e.** SOD1 protein levels varied significantly between mouse strains and were increased in vehicle-treated SOCK and h*SOD1^WT^*mice compared with wild-type and *Ctr1^+/−^* mice. CuATSM treatment elicited increases in SOD1 protein levels in wild-type and *Ctr1^+/−^*mice. Full representative immunoblots are presented in **Supplementary Figure 17**. Data in panels **a**, **b** and **e** represent mean ± SEM (*n* = 9-11/genotype/treatment), with full details of statistical tests presented in **Supplementary Table 2**. Comparisons marked with an asterisk (*) denote those made between vehicle- and CuATSM-treated mice of the same genotype, those marked with an arrowhead (^) demarcate those made to vehicle-treated wild-type mice, while those marked with a hashtag (#) denote those made to vehicle-treated h*SOD1^WT^* mice. ## *p* < 0.01, ### *p* < 0.001, \**p* < 0.05, \*\*\*\**p* < 0.0001, ^^^^*p* < 0.0001. **f.** SOD1 protein levels were positively correlated with total SOD activity in vehicle- and CuATSM-treated wild-type and *Ctr1^+/−^*mice. Statistical test details for **c**, **d** and **f** are presented in the panel.

### CuATSM rescues the nigrostriatal system in SOCK mice

The accumulation of disSOD1 pathology in the SOCK mouse SN is associated with significant changes in nigrostriatal neurotransmission from 6 months of age^13^. To determine whether decreasing levels of disSOD1 pathology in SOCK mice prevents these changes, we quantified levels of dopamine and its metabolite, homovanillic acid (HVA), as well as dopamine turnover (HVA normalized to dopamine), in the striatum of all four mouse strains following CuATSM and vehicle treatment using high performance liquid chromatography. Consistent with our previous findings in similar-aged animals^13^, vehicle-treated SOCK mice exhibited significantly lower levels of dopamine (**Fig. 5a**) and higher dopamine turnover (**Fig. 5b**) in the striatum compared with wild-type mice, with no changes identified in HVA levels between strains (**Supplementary Figure 18**). Dopamine levels were also lower in h*SOD1^WT^* mice, however dopamine turnover remained unchanged in this strain (**Fig. 5a,b**). Alterations to dopamine and dopamine turnover in SOCK mice were prevented by CuATSM treatment, to the extent that neither measure was significantly different between vehicle-treated wild-type mice and CuATSM-treated SOCK mice (**Fig. 5a,b**). By contrast, CuATSM treatment decreased striatal dopamine levels and increased dopamine turnover in h*SOD1^WT^* and wild-type mice.

**Figure 5.**
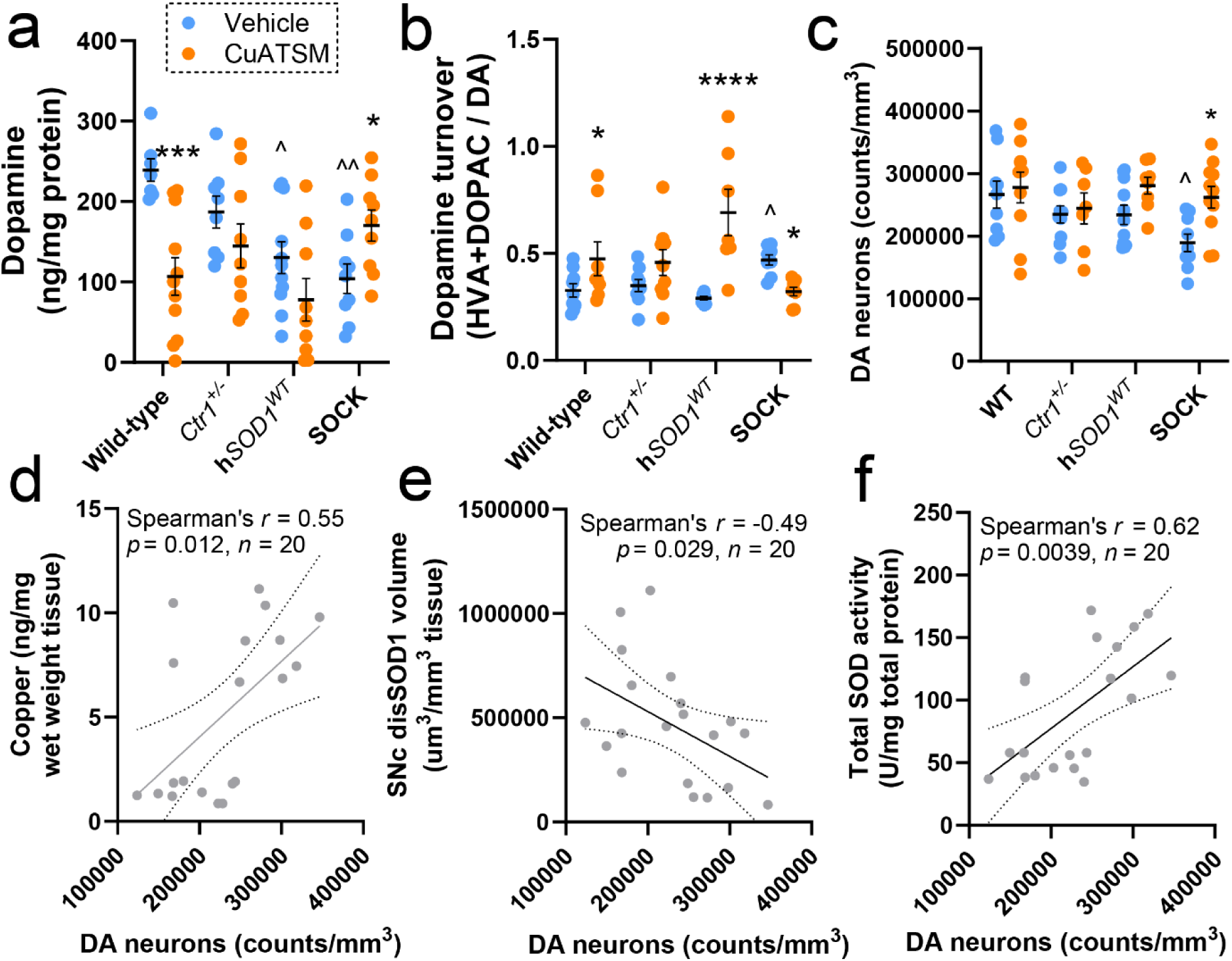
Evaluation of nigrostriatal neurotransmission and neurodegeneration in all mouse strains following treatment. **a.** Striatal dopamine levels were significantly reduced in h*SOD1^WT^*and SOCK mice treated with vehicle compared with wild-type mice, whilst CuATSM treatment increased striatal dopamine levels in SOCK mice and decreased them in wild-type mice. **b.** Striatal dopamine turnover, defined as the levels of homovanillic acid (HVA) and DOPAC normalized to dopamine (DA) levels, was significantly elevated in SOCK mice treated with vehicle compared with wild-type mice, with CuATSM treatment preventing this alteration and eliciting an elevation in dopamine turnover in wild-type and h*SOD1^WT^* mice. **c.** Stereological quantification of 3D-reconstructed tyrosine hydroxylase immunostaining revealed a significant loss of SNc dopamine neurons in vehicle-treated SOCK mice compared with all control mouse strains, which was rescued by CuATSM treatment. Representative 3D reconstructions of each genotype/treatment are presented in **Supplementary Figure 15**. Data in panels **a-c** represent mean ± SEM (*n* = 9-11/genotype/treatment), with full details of statistical tests presented in **Supplementary Table 2**. Comparisons marked with an asterisk (*) denote those made between vehicle- and CuATSM-treated mice of the same genotype, those marked with an arrowhead (^) demarcate those made to vehicle-treated wild-type mice. \**p* < 0.05, \*\*\**p* < 0.001, ^*p* < 0.05, ^^*p* < 0.01, ^^^*p* < 0.001. The density of SNc dopamine neurons was positively correlated with midbrain copper levels (**d**), inversely correlated with SNc disSOD1 volume (**e**) and positively correlated with midbrain total SOD activity (**f**) in SOCK mice. Statistical test details for **d - f** are presented in the panel.

Data from our previous study demonstrated that alterations to nigrostriatal neurotransmission are associated with dopamine neuron death in the SNc of SOCK mice^13^. To determine whether decreasing levels of disSOD1 pathology in SOCK mice decreases neuronal death, we performed stereological quantification of nigral dopamine neurons in all four mouse strains treated with vehicle and CuATSM using TH immunostaining of serial tissue sections spanning the entire SNc. Consistent with our previous findings in similar-aged animals^13^, vehicle-treated SOCK mice exhibited significant loss of SN dopamine neurons compared with wild-type (29% decrease) and h*SOD1^WT^* (19% decrease) mice, with no differences in the density of these neurons observed between all three control strains (**Fig. 5c**). CuATSM treatment prevented the death of these neurons in SOCK mice, maintaining their density at a similar level to that observed in vehicle-treated control strains (**Fig. 5c**). Normalization of striatal dopamine levels and turnover to SNc dopamine neuron density confirmed that CuATSM-induced improvements in striatal dopamine release and turnover in SOCK mice is likely a direct result of SNc dopamine neuron rescue (**Supplementary Figure 19**). Higher densities of SN dopamine neurons were also significantly correlated with higher tissue copper levels (**Fig. 5d**), lower disSOD1 volume (**Fig. 5e**) and higher total SOD activity (**Fig. 5f**) in the SNc of SOCK mice, suggesting that the observed neuroprotective effect of CuATSM may derive from the restoration of normal enzymatic activity and amelioration of the harmful effects of disSOD1.

### Treatment with CuATSM rescues motor impairment in SOCK mice

Prior examination of SOCK mouse motor performance using the rotarod test revealed robust motor impairment from 1.5 months-of-age^13^, which was observed in the absence of other features characterizing transgenic mutant SOD1 mice; weight loss, spinal motor neuron degeneration, and premature death^13^. To further characterize motor deficits in SOCK mice, we subjected vehicle-treated mice from all four strains to more comprehensive motor evaluation using balance beam, open field and grip strength tests. Vehicle-treated SOCK mice performed significantly poorer on the balance beam test compared with wild-type and h*SOD1^WT^* mice, taking 2.3-to-3.3-fold more time to cross the balance beam whilst exhibiting 11.2-to-19.2-fold greater paw slips (**Fig. 6a,b; Supplementary Video 1**). By contrast, all mouse strains possessed comparable forelimb grip strength (**Fig. 6c**), and no significant differences were observed for any parameters in the open field test, including time immobile, total distance travelled and central zone entries (**Fig. 6d-f**). Consistent with previous data^13^, there were also no differences in growth rates (**Fig. 6g**) or spinal motor neuron densities (**Fig. 6h**) between vehicle-treated mouse strains, demonstrating the absence of an overt ALS-like motor phenotype in SOCK mice.

**Figure 6.**
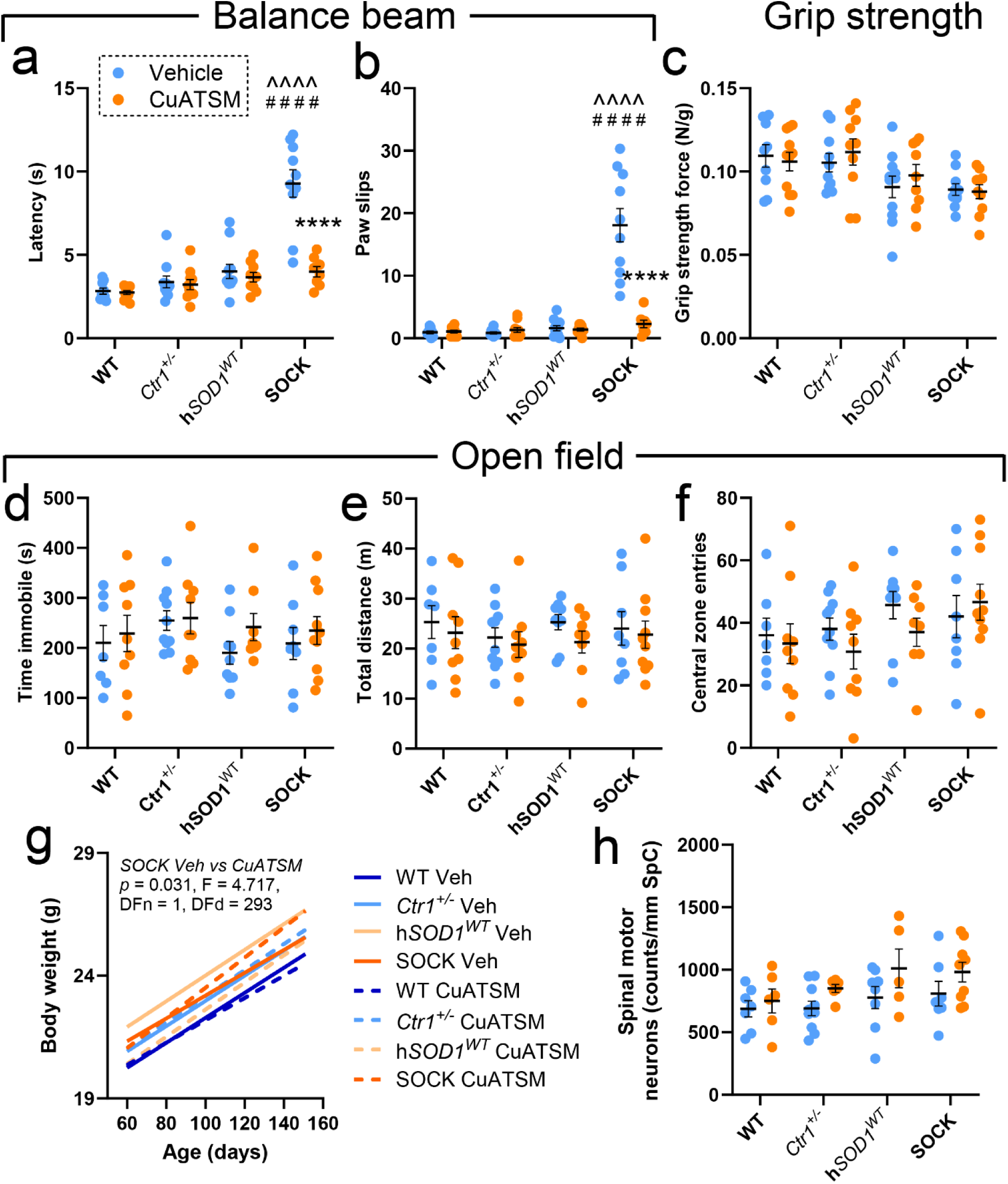
Motor performance testing in all mouse strains following treatment. In balance beam testing, the latency in time taken to cross the beam (**a**), together with the number of paw slips (**b**), varied significantly between mouse strains, with both increased in vehicle-treated SOCK mice compared with wild-type (WT) and h*SOD1^WT^* mice. CuATSM treatment significantly reduced both measures in SOCK mice, but did not change them in other strains. By contrast there was no variation in grip strength (**c**) or open field test performance metrics - time (sec) immobile (**d**), total distance travelled (meters; **e**), central zone entries (**f**) – between all strains and treatment groups. **g.** Mouse body weight increased at comparable rates between all four mouse strains treated with vehicle (*p* = 0.9093, F = 0.3890), although CuATSM treatment improved weight gain in SOCK mice (statistical test details presented in panel). **h.** Spinal motor neuron densities in each mouse genotype did not vary between vehicle and CuATSM treatment, nor did they vary between genotypes within the same treatment group. Data in panels **a-f** and **h** represent mean ± SEM (*n* = 9-11/genotype/treatment), with full details of statistical tests presented in **Supplementary Table 2**. Comparisons marked with an asterisk (*) denote those made between vehicle- and CuATSM-treated mice of the same genotype, those marked with an arrowhead (^) demarcate those made to vehicle-treated wild-type mice, while those marked with a hashtag (#) denote those made to vehicle-treated h*SOD1^WT^* mice. #### *p* < 0.0001, \*\*\*\**p* < 0.0001, ^^^^*p* < 0.0001.

Application of the same battery of behavioural tests to SOCK mice treated with CuATSM revealed striking rescue of motor impairment on the balance beam test (**Supplementary Video 2**), with no differences identified between these mice and vehicle-treated wild-type mice for this or any other motor test performed (**Fig. 6a-f**). Furthermore, CuATSM treatment improved weight gain in SOCK mice without negatively impacting the growth of other mouse strains (**Fig. 6g**), and did not alter the density of spinal motor neurons compared with vehicle in any mouse strain (**Fig. 6h**). These data highlight the ability of CuATSM at the current dosage to rescue motor impairment in SOCK mice without inducing clinical signs of toxicity, which have previously been reported following chronic CuATSM administration at higher doses^29^.

## Discussion

Previous work by our group demonstrated that reduced copper binding is likely a pivotal driver of wild-type disSOD1 pathology in post-mortem Parkinson disease brain tissues and SOCK mice^9, 13^. Copper is the key catalytic cofactor mediating SOD1 antioxidant activity and is also a vital structural cofactor, hence copper-deficient SOD1 is dysfunctional and exhibits greater structural flexibility^37^. This increases the solvent-exposure of amino acid residues normally buried within the protein and makes them more susceptible to atypical chemical modification^37, 38, 52^, which likely underlies many of the altered post-translational modifications that promote misfolding and aggregation of the protein in central nervous system tissues of Parkinson disease patients and SOCK mice^53^. We therefore proposed brain copper supplementation may improve SOD1 copper binding and mitigate the development of disSOD1 pathology in Parkinson disease. Data from the current study support this proposition, demonstrating a marked decrease in the formation of disSOD1 pathology, and higher SOD1 enzymatic activity, in SOCK mice following treatment with CuATSM, both of which were tightly correlated with tissue copper levels. While similar benefits have been reported in transgenic mutant SOD1 mouse models of ALS^14, 28, 54^, this is the first study to demonstrate similar benefits of this approach for wild-type SOD1 pathology *in vivo*.

Existing research into mutant and wild-type disSOD1 has largely focused on the accumulation of this pathology within vulnerable neuronal populations, especially spinal motor neurons, however mounting data implicates astrocytes in the development and propagation of this pathology^55–57^. Our study adds to this data by demonstrating that Parkinson-like wild-type disSOD1 accumulates to a greater extent within astrocytes than dopamine neuron soma in the SOCK mouse SN. Furthermore, we show that this appears to be driven by copper-dependent mechanisms, with CuATSM treatment decreasing disSOD1 within astrocytes to a much greater extent (8.5-fold) than within neurons (2-fold) or other compartments (3.8-fold). This is perhaps unsurprising given the pivotal role of this cell type in brain copper homeostasis, and may explain the expression of astrocytic SOD1 inclusions in transgenic mutant SOD1 mice^31^ and *SOD1*-linked ALS patient tissues^32^, both of which are characterized by altered copper homeostasis^58, 59^. Further targeted investigations into astrocytes as a nexus between copper and SOD1 pathology are warranted to determine whether astrocytes are particularly susceptible to developing this pathology *in vivo*, whether they play a role in its spread^55^, or whether they simply take up disSOD1 from dying neurons. For these investigations, it will be important to include additional astrocyte markers to complement GFAP immunoreactivity, given GFAP is not expressed by all astrocyte subtypes^60^. Beyond the specific role of astrocytes in the development of disSOD1 pathology, it is unclear which cell types and/or compartments contain the remaining ∼70% of disSOD1 pathology in the SN of SOCK mice. The very low volume of this pathology within SNc dopamine neuron soma suggests that disSOD1 deposition may instead occur in axons or dendrites, or may arise in alternative cell types such as oligodendrocytes or microglia, as observed in transgenic mutant SOD1 mice^61^ and ALS patients^32^, warranting further investigation.

In addition to advancing our understanding of mechanisms governing the formation of wild-type disSOD1 pathology, our data clearly demonstrates the positive impact of decreasing this pathology on SN dopamine neuron survival in SOCK mice. Structurally-disordered wild-type and mutant SOD1 are thought to contribute to neurodegeneration in Parkinson disease and ALS through simultaneous loss- and gain-of-protein functions^9, 37, 53, 62^. Here, disSOD1 is unable to perform its normal protective antioxidant function (loss), instead participating in atypical interactions that promote cellular damage (gain). One such example is a novel interaction with the anti-apoptotic mitochondrial protein Bcl-2 in protein extracts from transgenic mutant SOD1 mice and ALS patients, which can inhibit mitochondrial permeability, hyperpolarize mitochondria and lead to cell death^63, 64^. At present, it is difficult to ascertain which of these mechanisms may underlie disSOD1-mediated toxicity in SOCK mice, and which are corrected by CuATSM treatment to rescue nigral dopamine neurons. In particular, the significant increase in total SOD activity in vehicle-treated SOCK mice compared with non-transgenic control strains appears to challenge the existence of a SOD1 loss-of-function in these mice, however it must be acknowledged that there is no data on redox balance within the SOCK mouse SN. Given disSOD1 pathology promotes mitochondrial dysfunction and oxidative stress^65^, it is possible that superoxide is elevated to an even greater extent than SOD activity in SOCK mice, hence a SOD1 loss-of-function cannot be discounted. Further studies examining the lost and acquired function(s) of wild-type disSOD1, and the impact of these changes on nigral dopamine neurons, will be necessary to better understand how this pathology contributes to dopamine neuron loss in the Parkinson disease SN.

Aside from the question of SOD1 functionality, it is interesting that disSOD1 pathology exhibits such a clear relationship with dopamine neuron vulnerability in the SNc, despite the minimal presence of disSOD1 deposition within the soma of these neurons. We posit that disSOD1 may therefore impart toxicity to these neurons in a manner consistent with the dying back theory of neurodegeneration in Parkinson disease^66^, whereby the degeneration of SNc dopamine neuron projections precedes loss of their soma. Such an explanation is consistent with the observed decrease in dopamine levels and increase in dopamine turnover in the striatum, and may underlie the inverse correlation between SNc dopamine neuron density and disSOD1 volume in the SNr where these projects are located (**Supplementary Figure 16**), although disSOD1 has not yet been quantified in the striatum of SOCK mice. This is observed in other animal models of Parkinson disease, whereby the destruction of SN dopamine neuron terminals in the striatum occurs before that of cell bodies in monkeys treated with MPTP neurotoxin^67^, while protecting these terminals prevents the death of cell bodies following MPTP treatment in mice^68^. The mechanisms underlying this process remains an open question. Astrocytes and oligodendrocytes expressing mutant SOD1 how been shown to trigger the death of spinal motor neurons through the release of toxic factors^55, 61^. It is therefore possible that those containing wild-type disSOD1 in the SNr identified in this study may also contribute to the death of dopaminergic projections from the SNc. Further studies examining the exact subcellular localization of disSOD1 outside of dopamine neuron soma and astrocytes are warranted.

When extrapolating the findings of this study to Parkinson disease patients, it is important to consider that the genetic changes used to precipitate wild-type disSOD1 pathology in SOCK mice were introduced globally. This does not reflect the regional expression of disSOD1 in the post-mortem Parkinson disease brain^9^, which is largely restricted to degenerating regions such as the SNc, with disSOD1 deposition identified outside of these regions in SOCK mice^13^. This is particularly important to consider in relation to the robust motor phenotype exhibited by SOCK mice, which may be precipitated by disSOD1 within multiple brain regions important for movement, including the SNc but also other regions such as the cerebellum. These have not yet been examined as our primary aim was to interrogate the impact of brain copper supplementation on mechanisms governing the formation of wild-type disSOD1 pathology and if this could improve dopamine neuron viability in the SNc.

Following previous data from our group implicating wild-type disSOD1 accumulation in the death of nigral dopamine neurons, we now provide the first empirical evidence that reducing this pathology using copper supplementation can mitigate dopamine neuron loss. In considering how such an approach may best be translated into humans, it is important to recognize that the different genetic compositions of the mouse strains employed in this study significantly impacted the response to CuATSM. The increase in astrocyte volume in strains expressing normal levels of *Ctr1*, for example, may reflect astrogliosis from brain copper overload^69^, which when excessive or prolonged can lead to the formation of a glial scar that hinders neuronal homeostasis and the regeneration of damaged neurons^70^. This was likely avoided in SOCK and *Ctr1^+/−^* mice upon treatment with the same dose of CuATSM due to underlying brain copper deficiency in these strains. CuATSM also stimulated a 1.5-to-2.1-fold increase in SOD1 protein expression in strains lacking the h*SOD1^WT^*transgene, which we propose resulted from increased mitochondrial respiration and superoxide generation^71^. This did not occur in SOCK and h*SOD1^WT^* mice as SOD1 protein levels were already elevated 12.7-fold, which likely drew copper away from mitochondria in favour of nascent SOD1 protein^58^. While the extent of these differences is not expected to be as severe between patients, our data highlight the importance of titrating copper doses to individual patients, ensuring adequate copper supply is achieved without risking toxicity associated with excess brain copper. This is even more challenging in Parkinson disease patients given copper is only reduced in degenerating brain regions^12^, necessitating methods for targeting copper delivery to these regions. Importantly, these approaches are currently limited by the lack of a method for measuring brain copper levels *in vivo*.

Accompanying copper-dependent mechanisms of disSOD1 formation, data from this study indicates a small proportion of disSOD1 pathology may form via copper-independent mechanisms. Indeed, vehicle-treated h*SOD1^WT^* mice possessed low quantities of this pathology that were not significantly impacted by CuATSM treatment and were comparable in volume to the amount of disSOD1 remaining in SOCK mice following CuATSM treatment. These data align with previous data by our group demonstrating that mutant and wild-type SOD1 pathology in ALS patients likely forms from several convergent mechanisms^21^, suggesting that coadministration of additional therapies targeting these as-of-yet unknown mechanisms may be required to completely ameliorate Parkinson-like wild-type disSOD1 pathology. Given such combinational therapies often increase the incidence of adverse events^72^, it will first be important to determine the extent of reduction in this pathology that is required to achieve meaningful clinical outcomes and if this can be met using monotherapies against disSOD1 alone. Alternatively, these data may justify the evaluation of therapies that decrease SOD1 protein production as effective treatments against wild-type disSOD1 pathology in Parkinson disease, such as the FDA-approved antisense oligonucleotide Tofersen^73^. Indeed, increased SOD1 protein production constitutes a key factor driving the development of disSOD1 pathology in SOCK mice and Parkinson disease^9, 13^. Here the benefits of reducing the levels of such an important antioxidant protein must be carefully considered, with data from *Sod1* knock-out mice demonstrating that this can greatly increase susceptibility to neuron death following exposure to neurotoxic factors^62^, despite the absence of an overt neurological phenotype.

## Conclusions

This study provides the first *in vivo* evidence that copper supplementation can counteract the formation of Parkinson-like wild-type disSOD1 pathology and improve the survival of nigral dopamine neurons. Our data, combined with the presence of this pathology in idiopathic Parkinson disease, suggest that this pathology may represent a novel but treatable cell death pathway shared by the majority of patients. Importantly, it is likely that the positive impact of brain copper supplementation on disSOD1 pathology can be augmented by the co-administration of therapies which stabilize the protein via copper-independent mechanisms – an exciting avenue for future investigation.

## Abbreviations

ALS: amyotrophic lateral sclerosis
BSA: bovine serum albumin
CCS: copper chaperone for SOD1
CNS: central nervous system
Ct: control
Ctr1: copper transport protein 1
CuATSM: diacetyl-bis(4-methylthiosemicarbazonato)copper(II)
DIA: data independent acquisition
disSOD1: disordered SOD1
DOPAC: 3,4-dihydroxyphenylacetic acid
ER: endoplasmic reticulum
HPLC: high performance liquid chromatography
h*SOD1^WT^*: wild-type human SOD1 (gene)
HVA: homovanillic acid
ICP-MS: inductively coupled plasma mass spectrometry
PD: Parkinson disease
PTM: post-translational modification
SEM: standard error of the mean
SNc: substantia nigra pars compacta
SNr: substantia nigra pars reticulata
SOCK: SOD1-overexpressing Ctr1-knockdown (mice)
SOD1: Cu-Zn superoxide dismutase 1
SOD2: Mn superoxide dismutase
SOD3: extracellular superoxide dismutase
SpC: spinal cord
TH: tyrosine hydroxylase
UβB: unfolded beta barrel
WT: wild-type.

## Declarations

### Ethics approval and consent to participate

All experimental procedures involving the use of mice conformed to the Australian Code of Practice for the Care and Use of Animals for Scientific Purposes, with protocols approved by the Animal Ethics Committee at the University of Sydney (Ethics ID: 2020/1849).

### Consent for publication

All authors read and approved the final manuscript prior to publication.

### Availability of data and scripts

All associated data from this manuscript are available from the corresponding author upon reasonable request. All custom scripts used to collect data in this manuscript are publicly-available on GitHub (owner: Richard Harwood, repository: Sod1_CuATSM_Image_Analysis; https://github.com/RichardHarwood/Sod1_CuATSM_Image_Analysis).

### Author contributions

K.L.D. conceptualized, funded and supervised the study with input from B.D.R. and B.G.T. Animal ethics was obtained by B.D.R and K.L.D. CuATSM synthesis was performed by M.G. with assistance from R.C. Generation, treatment and behavioral analysis of mice was performed by B.D.R with assistance from C.K., S.R., V.C. and K.L.D. Tissue collection was conducted by B.D.R. with assistance from C.K., S.R. and K.L.D. Mouse tissue processing for fixed and fresh tissue analyses was performed by B.D.R., C.K. and B.G.T. Collection and analysis of histological data from fixed mouse tissues was performed by G.S., B.G.T, R.H., V.C., A.H.A., M.R. and A.A.L. Collection of biochemical data from fresh frozen mouse tissues was performed by B.G.T., D.M., B.D.R., V.C., F.P. and A.A, with analyses completed by B.G.T. Manuscript writing was completed by B.G.T. and K.L.D. with input from all authors.

## Acknowledgements

The authors acknowledge the facilities, as well as the scientific and technical assistance, of the Australian Microscopy and Microanalysis Research Facility (http://ammrf.org.au) node at the University of Sydney, as well as those of the Sydney Mass Spectrometry Facility at the University of Sydney.

