## Supplementary material for "Copper supplementation mitigates Parkinson-like wild-type SOD1 pathology and nigrostriatal degeneration in a novel mouse model"

### SUPPLEMENTARY INFORMATION

Supplementary Tables 1 - 4

1. Primary antibody details for all methods used in this study.
2. Statistical test details for all reported comparisons.
3. Volume of disSOD1 pathology localized within dopamine neurons and astrocytes, as well as independent of both cell types.
4. Altered SOD1 PTMs in the SN of vehicle- and CuATSM-treated SOCK mice compared with vehicle-treated h*SOD1*$^{WT}$ mice.

Supplementary Figures 1 - 19

1. $^{1}$H NMR Spectrum (600 MHz, DMSO-$d_6$) of $ATSMH_2$
2. $^{13}$C NMR Spectrum (125 MHz, DMSO-$d_6$) of $ATSMH_2$
3. HPLC-MS chromatogram of CuATSM after copper-loading and SPE method development
4. High-resolution mass spectrum of CuATSM
5. GAPDH protein levels in the midbrain of all mouse strains treated with CuATSM and vehicle.
6. Negative control immunofluorescent staining.
7. Summary of image processing and analysis workflow used to count dopamine neurons and measure the volume of disSOD1 and astrocytes in the SNc of all mouse strains.
8. Estimate of the accuracy of re-trained Cellpose deep learning model for segmenting and quantifying SNc dopamine neurons.
9. Immunofluorescent staining for spinal motor neurons in all four mouse genotypes following treatment.
10. Summary of behavioural test procedures performed on all mice.
11. Pilot study to determine CuATSM dose.
12. Zinc levels in the midbrain of all four mouse strains treated with vehicle or CuATSM.
13. Immunofluorescent characterization of disSOD1 pathology in the SNc of all four mouse strains treated with vehicle or CuATSM.
14. Immunofluorescent characterization of disSOD1 pathology in the SNr of all four mouse strains treated with vehicle or CuATSM.
15. Three dimensional reconstructions of disSOD1 pathology in the SNc and SNr of all mouse strains treated with vehicle and CuATSM.
16. Astrocyte volume in the SN of all mouse strains following vehicle and CuATSM treatment.

17. Representative SOD1 and GAPDH immunoblots from midbrain tissues of all four mouse genotypes.
18. Homovanillic acid levels in the striatum of all four mouse strains treated with vehicle or CuATSM
19. Striatal dopamine levels and turnover normalized to SNc dopamine neuron density in all four mouse strains treated with vehicle or CuATSM

**Supplementary Table 1. Primary antibody details for all methods used in this study.**

| Antibody | Source and catalogue # | Type | Host | Species reactivity | Immunogen | Application | Dilution |
|---|---|---|---|---|---|---|---|
| SOD1 UβB | StressMarq Biosciences, British Columbia, Canada (SPC-205) | P | Rb | Ms, Hu, Rat | N-terminal region, SOD1 protein with unfolded β-barrel | IF | 1:300 |
| Pan-SOD1 | Enzo Biochem, New York, USA (ADI-SOD-100) | P | Rb | Ms, Hu, Rat | Native human Cu/Zn SOD | WB<br>IP | 1:2000<br>12.5ug/mg beads |
| GAPDH | Sigma Aldrich, St Louis, Missouri, USA (G9545) | P | Rb | Ms, Hu, Rat | Synthetic peptide corresponding to amino acids of mouse GAPDH, conjugated to KLH via an N-terminal cysteine residue | WB | 1:1000 |
| TH | Abcam, Cambridge, United Kingdom (ab76442) | P | Ch | Ms | Synthetic peptides corresponding to sequences shared between murine (P24529) and human (P07101) TH | IF | 1:300 |
| GFAP | Thermo Fisher, Waltham, Massachusetts, USA (13-0300) | P | Rat | Ms, Hu, Rat | Enriched bovine glial filaments | IF | 1:10,000 |
| ISL1 | Abcam, Cambridge, United Kingdom (ab109517) | M | Rb | Ms, Hu, Rat | N/A | IF | 1:1000 |
| ChAT | Sigma Aldrich, St Louis, Missouri, USA (AB144P) | P | Gt | Ms, Hu, Rat | Human placental ChAT | IF | 1:500 |

**Abbreviations:** Ch, chicken; Gt, goat; Hu, human; IF, immunofluorescence; IP, immunoprecipitation; M, monoclonal; Ms, mouse; P, polyclonal; Rb, rabbit; SOD1, superoxide dismutase 1; TH, Tyrosine hydroxylase; UβB, unfolded beta-barrel.

**Supplementary Table 2. Statistical test details for all reported comparisons.** Statistically significant results in **bold**. Level of significance (α) = 0.05 for all comparisons.

| Display item | Parameter | Comparison groups | Test | Test statistics | Post-hoc test | Comparison groups (adjusted *p* value) |
|---|---|---|---|---|---|---|
| Fig. 1d | SN disSOD1 volume | All mouse strains, vehicle and CuATSM-treated | Two-way ANOVA | **Genotype: $F_{(3, 67)}$ = 33.69, $p < 0.0001$**<br><br>**Treatment: $F_{(1, 67)}$ = 29.87, $p < 0.0001$** | Dunnet's | WT vehicle vs CuATSM (>0.9999)<br>$Ctr1^{+/-}$ vehicle vs CuATSM (0.8216)<br>h$SOD1^{WT}$ vehicle vs CuATSM (0.6716)<br>**SOCK vehicle vs CuATSM (<0.0001)**<br>Vehicle: WT vs $Ctr1^{+/-}$ (0.5684)<br>**Vehicle: WT vs h$SOD1^{WT}$ (0.0029)**<br>**Vehicle: WT vs SOCK (<0.0001)**<br>**Vehicle: h$SOD1^{WT}$ vs SOCK (<0.0001)**<br>CuATSM: WT vs $Ctr1^{+/-}$ (0.9903)<br>CuATSM: WT vs h$SOD1^{WT}$ (0.1004)<br>CuATSM: WT vs SOCK (0.1474)<br>CuATSM: h$SOD1^{WT}$ vs SOCK (0.9840) |
| Fig. 1e | Midbrain copper levels | All mouse strains, vehicle and CuATSM-treated | Two-way ANOVA | **Genotype: $F_{(3, 72)}$ = 19.17, $p < 0.0001$**<br><br>**Treatment: $F_{(1, 72)}$ = 246.0, $p < 0.0001$** | Dunnet's | **WT vehicle vs CuATSM (<0.0001)**<br>**$Ctr1^{+/-}$ vehicle vs CuATSM (<0.0001)**<br>**h$SOD1^{WT}$ vehicle vs CuATSM (<0.0001)**<br>**SOCK vehicle vs CuATSM (<0.0001)**<br>**Vehicle: WT vs $Ctr1^{+/-}$ (0.0030)**<br>Vehicle: WT vs h$SOD1^{WT}$ (0.9767)<br>**Vehicle: WT vs SOCK (0.0252)**<br>**Vehicle: h$SOD1^{WT}$ vs SOCK (0.0075)**<br>**CuATSM: WT vs $Ctr1^{+/-}$ (<0.0001)**<br>CuATSM: WT vs h$SOD1^{WT}$ (0.8812)<br>CuATSM: WT vs SOCK (0.4371)<br>CuATSM: h$SOD1^{WT}$ vs SOCK (0.8931) |
| Fig. 4a | Total SOD activity | All mouse strains, vehicle and CuATSM-treated | Two-way ANOVA | **Genotype: $F_{(3, 72)}$ = 268.3, $p < 0.0001$**<br><br>**Treatment: $F_{(1, 72)}$ = 178.2, $p < 0.0001$** | Dunnet's | **WT vehicle vs CuATSM (0.0232)**<br>**$Ctr1^{+/-}$ vehicle vs CuATSM (<0.0001)**<br>**h$SOD1^{WT}$ vehicle vs CuATSM (<0.0001)**<br>**SOCK vehicle vs CuATSM (<0.0001)**<br>Vehicle: WT vs $Ctr1^{+/-}$ (0.9981)<br>**Vehicle: WT vs h$SOD1^{WT}$ (<0.0001)**<br>**Vehicle: WT vs SOCK (<0.0001)**<br>**Vehicle: h$SOD1^{WT}$ vs SOCK (0.0003)**<br>**CuATSM: WT vs $Ctr1^{+/-}$ (0.0069)**<br>**CuATSM: WT vs h$SOD1^{WT}$ (<0.0001)** |

| | | | | | | |
|---|---|---|---|---|---|---|
| | | | | | | **CuATSM: WT vs SOCK (<0.0001)**<br>CuATSM: h*SOD1*$^{WT}$ vs SOCK (0.9989) |
| Fig. 4b | SOD activity per unit of SOD1 protein | All mouse strains, vehicle and CuATSM-treated | Two-way ANOVA | **Genotype: $F_{(3, 70)}$ = 35.07, $p$ < 0.0001**<br><br>**Treatment: $F_{(1, 70)}$ = 34.19, $p$ < 0.0001** | Dunnet's | WT vehicle vs CuATSM (0.7185)<br>*Ctr1*$^{+/-}$ vehicle vs CuATSM (0.8465)<br>**h*SOD1*$^{WT}$ vehicle vs CuATSM (<0.0001)**<br>**SOCK vehicle vs CuATSM (<0.0001)**<br>Vehicle: WT vs *Ctr1*$^{+/-}$ (0.4437)<br>**Vehicle: WT vs h*SOD1*$^{WT}$ (<0.0001)**<br>**Vehicle: WT vs SOCK (<0.0001)**<br>**Vehicle: h*SOD1*$^{WT}$ vs SOCK (0.0044)**<br>CuATSM: WT vs *Ctr1*$^{+/-}$ (0.5257)<br>CuATSM: WT vs h*SOD1*$^{WT}$ (0.4216)<br>CuATSM: WT vs SOCK (0.3158)<br>CuATSM: h*SOD1*$^{WT}$ vs SOCK (0.9990) |
| Fig. 4e | SOD1 protein levels | All mouse strains, vehicle and CuATSM-treated | Two-way ANOVA | **Genotype: $F_{(3, 70)}$ = 312.3, $p$ < 0.0001**<br><br>**Treatment: $F_{(1, 70)}$ = 9.078, $p$ = 0.0036** | Dunnet's | **WT vehicle vs CuATSM (0.0071)**<br>***Ctr1*$^{+/-}$ vehicle vs CuATSM (<0.0001)**<br>h*SOD1*$^{WT}$ vehicle vs CuATSM (0.4272)<br>SOCK vehicle vs CuATSM (0.5803)<br>Vehicle: WT vs *Ctr1*$^{+/-}$ (0.3987)<br>**Vehicle: WT vs h*SOD1*$^{WT}$ (<0.0001)**<br>**Vehicle: WT vs SOCK (<0.0001)**<br>Vehicle: h*SOD1*$^{WT}$ vs SOCK (0.9997)<br>**CuATSM: WT vs *Ctr1*$^{+/-}$ (0.0004)**<br>**CuATSM: WT vs h*SOD1*$^{WT}$ (<0.0001)**<br>**CuATSM: WT vs SOCK (<0.0001)**<br>CuATSM: h*SOD1*$^{WT}$ vs SOCK (0.9929) |
| Fig. 5a | Striatal dopamine levels | All mouse strains, vehicle and CuATSM-treated | Two-way ANOVA | **Genotype: $F_{(3, 65)}$ = 4.102, $p$ = 0.0100**<br><br>**Treatment: $F_{(1, 65)}$ = 6.502, $p$ = 0.0131** | Tukey's | **WT vehicle vs CuATSM (0.0001)**<br>*Ctr1*$^{+/-}$ vehicle vs CuATSM (0.1978)<br>h*SOD1*$^{WT}$ vehicle vs CuATSM (0.0845)<br>**SOCK vehicle vs CuATSM (0.0398)**<br>Vehicle: WT vs *Ctr1*$^{+/-}$ (0.4388)<br>**Vehicle: WT vs h*SOD1*$^{WT}$ (0.0068)**<br>**Vehicle: WT vs SOCK (0.0009)**<br>Vehicle: h*SOD1*$^{WT}$ vs SOCK (0.8156)<br>CuATSM: WT vs *Ctr1*$^{+/-}$ (0.5891)<br>CuATSM: WT vs h*SOD1*$^{WT}$ (0.7680)<br>CuATSM: WT vs SOCK (0.1623)<br>**CuATSM: h*SOD1*$^{WT}$ vs SOCK (0.0236)** |

| | | | | | | |
|---|---|---|---|---|---|---|
| Fig. 5b | Striatal dopamine turnover | All mouse strains, vehicle and CuATSM-treated | Two-way ANOVA | Genotype: $F_{(3, 59)} = 0.7728$, $p = 0.5138$<br><br>**Treatment: $F_{(1, 59)} = 10.38$, $p = 0.0021$** | Tukey's | **WT vehicle vs CuATSM (0.0315)**<br>*$Ctr1^{+/-}$* vehicle vs CuATSM (0.1211)<br>**h*$SOD1^{WT}$* vehicle vs CuATSM (<0.0001)**<br>**SOCK vehicle vs CuATSM (0.0137)**<br>Vehicle: WT vs *$Ctr1^{+/-}$* (0.9570)<br>Vehicle: WT vs h*$SOD1^{WT}$* (0.9307)<br>**Vehicle: WT vs SOCK (0.0415)**<br>**Vehicle: h*$SOD1^{WT}$* vs SOCK (0.0110)**<br>CuATSM: WT vs *$Ctr1^{+/-}$* (0.9977)<br>CuATSM: WT vs h*$SOD1^{WT}$* (0.0803)<br>CuATSM: WT vs SOCK (0.1577)<br>**CuATSM: h*$SOD1^{WT}$* vs SOCK (0.0002)** |
| Fig. 5c | SNc dopamine neuron density | All mouse strains, vehicle and CuATSM-treated | Two-way ANOVA | Genotype: $F_{(3, 67)} = 2.471$, $p = 0.0693$<br><br>**Treatment: $F_{(1, 67)} = 7.079$, $p = 0.0098$** | Dunnet's | WT vehicle vs CuATSM (0.9871)<br>*$Ctr1^{+/-}$* vehicle vs CuATSM (0.9948)<br>h*$SOD1^{WT}$* vehicle vs CuATSM (0.2847)<br>**SOCK vehicle vs CuATSM (0.0220)**<br>Vehicle: WT vs *$Ctr1^{+/-}$* (0.8332)<br>Vehicle: WT vs h*$SOD1^{WT}$* (0.8021)<br>**Vehicle: WT vs SOCK (0.0351)**<br>Vehicle: h*$SOD1^{WT}$* vs SOCK (0.1947)<br>CuATSM: WT vs *$Ctr1^{+/-}$* (0.5485)<br>CuATSM: WT vs h*$SOD1^{WT}$* (0.9994)<br>CuATSM: WT vs SOCK (0.8938)<br>CuATSM: h*$SOD1^{WT}$* vs SOCK (0.8316) |
| Fig. 6a | Balance beam (latency) | All mouse strains, vehicle and CuATSM-treated | Two-way ANOVA | **Genotype: $F_{(3, 69)} = 31.98$, $p < 0.0001$**<br><br>**Treatment: $F_{(1, 69)} = 24.49$, $p < 0.0001$** | Dunnet's | WT vehicle vs CuATSM (>0.9999)<br>*$Ctr1^{+/-}$* vehicle vs CuATSM (>0.9999)<br>h*$SOD1^{WT}$* vehicle vs CuATSM (>0.9999)<br>**SOCK vehicle vs CuATSM (<0.0001)**<br>Vehicle: WT vs *$Ctr1^{+/-}$* (>0.9999)<br>Vehicle: WT vs h*$SOD1^{WT}$* (0.7354)<br>**Vehicle: WT vs SOCK (<0.0001)**<br>**Vehicle: h*$SOD1^{WT}$* vs SOCK (<0.0001)**<br>CuATSM: WT vs *$Ctr1^{+/-}$* (>0.9999)<br>CuATSM: WT vs h*$SOD1^{WT}$* (0.9786)<br>CuATSM: WT vs SOCK (0.7356)<br>CuATSM: h*$SOD1^{WT}$* vs SOCK (>0.9999) |
| Fig. 6b | Balance beam (paw slips) | All mouse strains, vehicle and CuATSM-treated | Two-way ANOVA | **Genotype: $F_{(3, 70)} = 36.21$, $p < 0.0001$** | Dunnet's | WT vehicle vs CuATSM (>0.9999)<br>*$Ctr1^{+/-}$* vehicle vs CuATSM (>0.9999)<br>h*$SOD1^{WT}$* vehicle vs CuATSM (>0.9999) |

| | | | | | | |
|---|---|---|---|---|---|---|
| | | | | **Treatment: $F_{(1, 70)} = 28.07$, $p < 0.0001$** | | **SOCK vehicle vs CuATSM (<0.0001)**<br>Vehicle: WT vs *Ctr1*$^{+/-}$ (>0.9999)<br>Vehicle: WT vs h*SOD1*$^{WT}$ (>0.9999)<br>**Vehicle: WT vs SOCK (<0.0001)**<br>**Vehicle: h*SOD1*$^{WT}$ vs SOCK (<0.0001)**<br>CuATSM: WT vs *Ctr1*$^{+/-}$ (>0.9999)<br>CuATSM: WT vs h*SOD1*$^{WT}$ (>0.9999)<br>CuATSM: WT vs SOCK (>0.9999)<br>CuATSM: h*SOD1*$^{WT}$ vs SOCK (>0.9999) |
| Fig. 6c | Grip strength | All mouse strains, vehicle and CuATSM-treated | Two-way ANOVA | **Genotype: $F_{(3, 72)} = 5.602$, $p = 0.0016$**<br><br>Treatment: $F_{(1, 72)} = 0.2613$, $p = 0.6108$ | Dunnet's | Vehicle: WT vs *Ctr1*$^{+/-}$ (>0.9999)<br>Vehicle: WT vs h*SOD1*$^{WT}$ (0.5676)<br>Vehicle: WT vs SOCK (0.4563)<br>Vehicle: h*SOD1*$^{WT}$ vs SOCK (>0.9999)<br>CuATSM: WT vs *Ctr1*$^{+/-}$ (>0.9999)<br>CuATSM: WT vs h*SOD1*$^{WT}$ (>0.9999)<br>CuATSM: WT vs SOCK (0.5779)<br>CuATSM: h*SOD1*$^{WT}$ vs SOCK (0.9998) |
| Fig. 6d | Open field (time immobile) | All mouse strains, vehicle and CuATSM-treated | Two-way ANOVA | Genotype: $F_{(3, 62)} = 0.9162$, $p = 0.4384$<br><br>Treatment: $F_{(1, 62)} = 1.506$, $p = 0.2245$ | N/A | N/A |
| Fig. 6e | Open field (total distance) | All mouse strains, vehicle and CuATSM-treated | Two-way ANOVA | Genotype: $F_{(3, 62)} = 0.3898$, $p = 0.7608$<br><br>Treatment: $F_{(1, 62)} = 1.368$, $p = 0.2467$ | N/A | N/A |
| Fig. 6f | Open field (central zone entries) | All mouse strains, vehicle and CuATSM-treated | Two-way ANOVA | Genotype: $F_{(3, 62)} = 1.693$, $p = 0.1777$<br><br>Treatment: $F_{(1, 62)} = 0.8323$, $p = 0.3651$ | N/A | N/A |
| Fig. 6h | Spinal motor neurons | All mouse strains, vehicle and CuATSM-treated | Two-way ANOVA | Genotype: $F_{(3, 50)} = 2.139$, $p = 0.1071$<br><br>**Treatment: $F_{(1, 50)} = 6.718$, $p = 0.0125$** | Dunnet's | WT vehicle vs CuATSM (0.9786)<br>*Ctr1*$^{+/-}$ vehicle vs CuATSM (0.5391)<br>h*SOD1*$^{WT}$ vehicle vs CuATSM (0.2707)<br>SOCK vehicle vs CuATSM (0.4419) |

| | | | | | | |
|---|---|---|---|---|---|---|
| Supp Fig. 1 | Midbrain copper levels | Vehicle vs 15 vs 30 mg/kg CuATSM | One-way ANOVA | **$F_{(2, 23)} = 131.7$, $p < 0.0001$** | Dunnett's | **Vehicle vs 15mg/kg (<0.0001)**<br>**Vehicle vs 30mg/kg (<0.0001)** |
| Supp Fig. 2 | Midbrain zinc levels | All mouse strains, vehicle and CuATSM-treated | Two-way ANOVA | **Genotype: $F_{(3, 72)} = 55.06$, $p < 0.0001$**<br><br>Treatment: $F_{(1, 72)} = 0.0035$, $p = 0.9529$ | Dunnet's | WT vehicle vs CuATSM (0.6021)<br>*Ctr1*$^{+/-}$ vehicle vs CuATSM (>0.9999)<br>h*SOD1*$^{WT}$ vehicle vs CuATSM (>0.9999)<br>SOCK vehicle vs CuATSM (0.9998)<br>Vehicle: WT vs *Ctr1*$^{+/-}$ (0.3088)<br>**Vehicle: WT vs h*SOD1*$^{WT}$ (0.0002)**<br>**Vehicle: WT vs SOCK (0.0203)**<br>Vehicle: h*SOD1*$^{WT}$ vs SOCK (0.9993)<br>CuATSM: WT vs *Ctr1*$^{+/-}$ (>0.9999)<br>**CuATSM: WT vs h*SOD1*$^{WT}$ (<0.0001)**<br>**CuATSM: WT vs SOCK (<0.0001)**<br>CuATSM: h*SOD1*$^{WT}$ vs SOCK (>0.9999) |
| Supp Fig. 6a | SN astrocyte volume | All mouse strains, vehicle and CuATSM-treated | Two-way ANOVA | Genotype: $F_{(3, 63)} = 1.820$, $p = 0.1526$<br><br>**Treatment: $F_{(1, 63)} = 20.940$, $p < 0.0001$** | Dunnett's | **WT vehicle vs CuATSM (< 0.0001)**<br>*Ctr1*$^{+/-}$ vehicle vs CuATSM (0.3776)<br>**h*SOD1*$^{WT}$ vehicle vs CuATSM (0.0034)**<br>SOCK vehicle vs CuATSM (0.1188)<br>Vehicle: WT vs *Ctr1*$^{+/-}$ (0.9989)<br>Vehicle: WT vs h*SOD1*$^{WT}$ (>0.9999)<br>**Vehicle: WT vs SOCK (0.0201)**<br>**Vehicle: h*SOD1*$^{WT}$ vs SOCK (0.0148)** |
| Supp Fig. 8 | GAPDH protein levels | All mouse strains, vehicle and CuATSM-treated | Two-way ANOVA | Genotype: $F_{(3, 70)} = 1.609$, $p = 0.1951$<br><br>Treatment: $F_{(1, 70)} = 1.540$, $p = 0.2187$ | N/A | N/A |
| Supp Fig. 14 | HVA levels | All mouse strains, vehicle and CuATSM-treated | Two-way ANOVA | Genotype: $F_{(3, 56)} = 2.566$, $p = 0.0636$<br><br>Treatment: $F_{(1, 56)} = 0.08831$, $p = 0.7674$ | N/A | N/A |

| | | | | | | |
|---|---|---|---|---|---|---|
| Supp Fig. 19 | Dopamine levels normalized to SNc dopamine neuron density | All mouse strains, vehicle and CuATSM-treated | Two-way ANOVA | **Genotype: $F_{(3, 66)}$ = 3.856, *p* = 0.0132**<br><br>Treatment: $F_{(1, 66)}$ = 3.403, *p* = 0.0696 | Tukey's | **WT vehicle vs CuATSM (0.0113)**<br>*Ctr1*$^{+/-}$ vehicle vs CuATSM (0.9771)<br>h*SOD1*$^{WT}$ vehicle vs CuATSM (0.1172)<br>SOCK vehicle vs CuATSM (0.6821)<br>Vehicle: WT vs *Ctr1*$^{+/-}$ (0.9965)<br>Vehicle: WT vs h*SOD1*$^{WT}$ (0.4879)<br>Vehicle: WT vs SOCK (0.3113)<br>Vehicle: h*SOD1*$^{WT}$ vs SOCK (0.9782)<br>**CuATSM: WT vs *Ctr1*$^{+/-}$ (0.0474)**<br>CuATSM: WT vs h*SOD1*$^{WT}$ (0.9808)<br>CuATSM: WT vs SOCK (0.5884)<br>CuATSM: h*SOD1*$^{WT}$ vs SOCK (0.3997) |
| Supp Fig. 19 | Dopamine turnover normalized to SNc dopamine neuron density | All mouse strains, vehicle and CuATSM-treated | Two-way ANOVA | Genotype: $F_{(3, 56)}$ = 1.759, *p* = 0.1654<br><br>Treatment: $F_{(1, 56)}$ = 0.01119, *p* = 0.9161<br><br>**Interaction: $F_{(3, 56)}$ = 7.475, *p* = 0.0003** | Tukey's | WT vehicle vs CuATSM (0.3871)<br>*Ctr1*$^{+/-}$ vehicle vs CuATSM (0.6282)<br>**h*SOD1*$^{WT}$ vehicle vs CuATSM (0.0125)**<br>**SOCK vehicle vs CuATSM (0.0003)**<br>Vehicle: WT vs *Ctr1*$^{+/-}$ (0.7561)<br>Vehicle: WT vs h*SOD1*$^{WT}$ (0.9889)<br>**Vehicle: WT vs SOCK (0.0012)**<br>**Vehicle: h*SOD1*$^{WT}$ vs SOCK (0.0032)**<br>CuATSM: WT vs *Ctr1*$^{+/-}$ (0.9443)<br>CuATSM: WT vs h*SOD1*$^{WT}$ (0.1969)<br>CuATSM: WT vs SOCK (0.8288)<br>**CuATSM: h*SOD1*$^{WT}$ vs SOCK (0.0344)** |

**Abbreviations:** ANOVA, analysis of variance; SN, substantia nigra; WT, wild-type.

**Supplementary Table 3. Volume of disSOD1 pathology localized within dopamine neurons and astrocytes, as well as independent of both cell types.** Data represent mean (SD, *n*).

| | **Wild-type** | | ***Ctr1*$^{+/-}$** | | **h*SOD1*$^{WT}$** | | **SOCK** | |
|---|---|---|---|---|---|---|---|---|
| | **Vehicle** | **CuATSM** | **Vehicle** | **CuATSM** | **Vehicle** | **CuATSM** | **Vehicle** | **CuATSM** |
| **SNc DA** | 1166 (1609, 9) | 1146 (1028, 10) | 3168 (2497, 10) | 3082 (1960, 7) | 7635 (6506, 11) | 7480 (5168, 8) | 21128 (8606, 9) | 8907 (5826, 11) |
| **SNc ast** | 5570 (7111, 9) | 8403 (7147, 10) | 11936 (5893, 10) | 11802 (9807, 7) | 19609 (16366, 11) | 20645 (14069, 7) | 64791 (34389, 9) | 23571 (10843, 11) |
| **SNc oth** | 44062 (49391, 9) | 38115 (30447, 10) | 119218 (72482, 10) | 72462 (32298, 7) | 173735 (129438, 11) | 226986 (135476, 8) | 605403 (236014, 9) | 252390 (144765, 11) |
| **SNc** | 50798 (57188, 9) | 47664 (37913, 10) | 134322 (77768, 10) | 87347 (41991, 7) | 200978 (147724, 11) | 267498 (155113, 8) | 691321 (249722, 9) | 284869 (156767, 11) |
| **SNr DA** | 13 (34, 9) | 7 (17, 10) | 18 (52, 10) | 1 (4, 7) | 15 (32, 11) | 2 (3, 8) | 133 (166, 9) | 57 (129, 11) |
| **SNr ast** | 12205 (17759, 9) | 12006 (11898, 10) | 26644 (20709, 10) | 6138 (4307, 6) | 39325 (45042, 10) | 33127 (42732, 8) | 256053 (98104, 9) | 30506 (20255, 11) |
| **SNr oth** | 23262 (29561, 9) | 19921 (17978, 10) | 70873 (56608, 10) | 24332 (22686, 7) | 169105 (175475, 11) | 136326 (119111, 8) | 654478 (343899, 9) | 88083 (51687, 11) |
| **SNr** | 35480 (47154, 9) | 31935 (28915, 10) | 97536 (70857, 10) | 42359 (53838, 7) | 242355 (280988, 11) | 169455 (150303, 8) | 910664 (395596, 9) | 118646 (69485, 11) |

**Abbreviations:** ast, astrocytes; DA, dopamine neuron soma; oth, other; SNc, substantia nigra pars compacta; SNr, substantia nigra pars reticulata.

**Supplementary Table 4. Altered SOD1 PTMs in the SN of vehicle- and CuATSM-treated SOCK mice compared with vehicle-treated h*SOD1*$^{WT}$ mice.** SOD1 protein was isolated from post-mortem tissues using our published immunoprecipitation method [1], which we have demonstrated does not significantly alter SOD1 PTMs. Peptides were also prepared as previously described [1] and bottom-up proteomic mass spectrometry performed on a Q Exactive HF-X Hybrid Quadrupole-Orbitrap Mass Spectrometer operated in data-independent acquisition mode. Data were analyzed using Spectronaut proteomics software (Version 18, Biognosys, Schlieren, Zurich, Switzerland). Full methods details are presented in the main methods section. Amino acid residues are identified using one letter code. Sample size = 10/group. Atypical PTMs (top) are separated from physiological PTMs (bottom).

| Modification | Altered vehicle-treated SOCK mice vs vehicle-treated h*SOD1*$^{WT}$ mice ($Log_2$(ratio)) | Test statistics (p, Q, # ratios) | Alteration present in CuATSM-treated SOCK mice? ($Log_2$(ratio)) | Test statistics (p, Q, # ratios) |
|---|---|---|---|---|
| Oxidation (H) | H120 (1.12) | 0.000518619, 0.001555857, 84 | No | N/A |
| | H110 (1.00) | 0.027045034, 0.029503674, 82 | No | N/A |
| | H46 (0.97) | 0.010237982, 0.0124946, 82 | No | N/A |
| | H43 (0.97) | 0.010412167, 0.0124946, 82 | No | N/A |
| | H48 (2.83) | 1.20E-09, 1.43E-08, 80 | No | N/A |
| | H63 (3.57) | 0.00485058, 0.00970116, 10 | Yes (1.07) | 0.029366508, 0.004894418, 36 |
| Oxidation (W) | No | N/A | N/A | N/A |
| Kynurenine (W) | No | N/A | N/A | N/A |
| Hydroxykynurenine (W) | No | N/A | N/A | N/A |
| Dioxidation (W) | No | N/A | N/A | N/A |
| Nitration (W) | No | N/A | N/A | N/A |
| Carboxymethyllysine (K) | No | N/A | N/A | N/A |
| Glycation (R, K) | R79 (0.84) | 1.38E-16, 6.91E-16, 28 | No | N/A |
| | K30 (3.80) | 6.00E-06, 1.00E-05, 20 | Yes (0.85) | 6.28E-13, 1.13E-11, 80 |
| Gloxal AGE (R) | No | N/A | N/A | N/A |
| Acetylation (K) | K9 (-0.85) | 3.66E-07, 3.66E-07, 84 | No | N/A |
| | K128 (-1.21) | 0.001108004, 0.000369335, 12 | Yes (-1.55) | 0.000638268, 0.002553071, 10 |
| | K30 (-0.81) | 4.48E-06, 2.24E-06, 76 | No | N/A |

| | | | | |
|---|---|---|---|---|
| Succinylation (K) | K91 (0.91) | 4.05E-07, 1.35E-06, 84 | Yes (1.42) | 1.92E-08, 2.30E-07, 84 |
| | K9 (-0.69) | 8.85E-08, 4.42E-07, 84 | No | N/A |
| | K136 (-0.82) | 0.037085879, 0.037085879, 18 | No | N/A |
| | K3 (-0.76) | 0.010460055, 0.017433424, 32 | No | N/A |
| Phosphorylation (S, T) | S25 (-4.22) | 0.000329629, 0.000109876, 80 | Yes (-2.64) | 0.018831555, 0.026902222, 82 |
| | S102 (-1.43) | 0.000487882, 0.000139395, 40 | No | N/A |
| | S134 (-0.70) | 0.002253022, 0.000500672, 84 | No | N/A |
| | T39 (-0.59) | 1.12E-05, 7.43E-06, 74 | No | N/A |
| | T88 (1.05) | 0.000702611, 0.000175653, 28 | No | N/A |
| Deamidation (N, Q) | N26 (-0.79) | 5.65E-08, 1.13E-07, 84 | No | N/A |
| | Q153 (-0.73) | 0.00114759, 0.000163941, 84 | No | N/A |
| | Q22 (1.45) | 0.001441793, 0.000192239, 46 | Yes (1.44) | 0.014949981, 0.025628539, 40 |
| | N139 (2.24) | 0.010717647, 0.00119085, 46 | Yes (1.28) | 0.01103089, 0.022061779, 40 |
| Ubiquitylation (GlyGly footprint; K) | K122 (-0.77) | 0.000909531, 0.001515885, 78 | No | N/A |
| | K91 (-0.78) | 0.000342425, 0.000972547, 84 | Yes (-2.69) | 0.008131769, 0.011616812, 14 |
| | K30 (-3.80) | 0.000571577, 0.001143154, 14 | Yes (-4.54) | 0.000265919, 0.001788504, 10 |
| Glycosylation (N, S, T) | S107 (-0.84) | 0.029912767, 0.034110507, 84 | No | N/A |
| | N131 (-1.70) | 0.015254411, 0.030508822, 84 | No | N/A |
| | S98 (-0.71) | 3.08E-06, 1.64E-05, 84 | No | N/A |
| | S102 (-0.95) | 0.030159266, 0.034110507, 16 | No | N/A |
| | N65 (-1.48) | 0.00050494, 0.001435579, 18 | Yes (-0.78) | 0.047831944, 0.01366627, 24 |
| | S34 (-1.47) | 0.00065419, 0.001495292, 36 | No | N/A |
| | T39 (0.99) | 0.000538342, 0.001435579, 18 | Yes (1.17) | 0.000387641, 0.000387641, 14 |
| Acetylglucosamine (N, S, T) | N131 (-0.71) | 2.17E-06, 5.50E-06, 84 | No | N/A |
| | T58 (-1.13) | 1.33E-06, 5.30E-06, 48 | No | N/A |
| | N26 (-1.42) | 0.000129834, 0.000207735, 14 | No | N/A |
| | S102 (1.06) | 0.034072718, 0.019470124, 10 | No | N/A |
| | S25 (-1.13) | 0.000224388, 0.000224388, 14 | No | N/A |
| | N19 (-1.13) | 0.000224388, 0.000224388, 14 | No | N/A |

**Abbreviations:** AGE, advanced glycation end-product; N/A, not applicable; SN, substantia nigra.

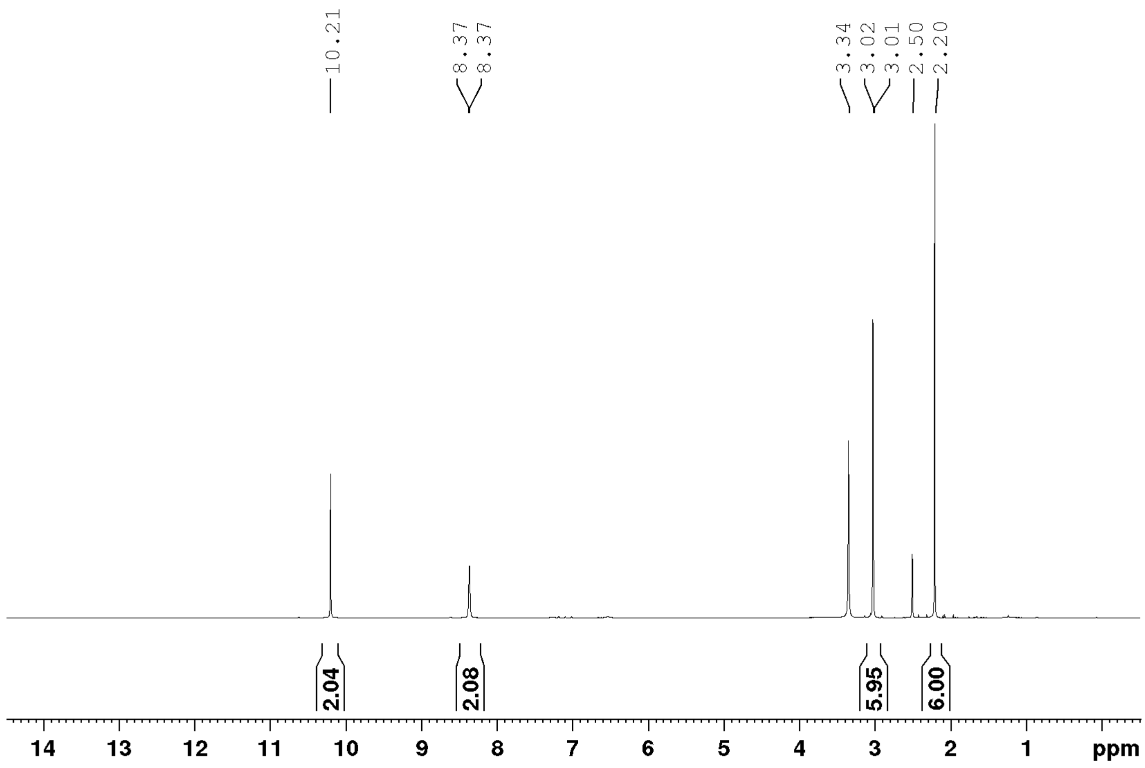


**Supplementary Figure 1. $^1$H NMR Spectrum (600 MHz, DMSO-$d_6$) of ATSMH$_2$.**

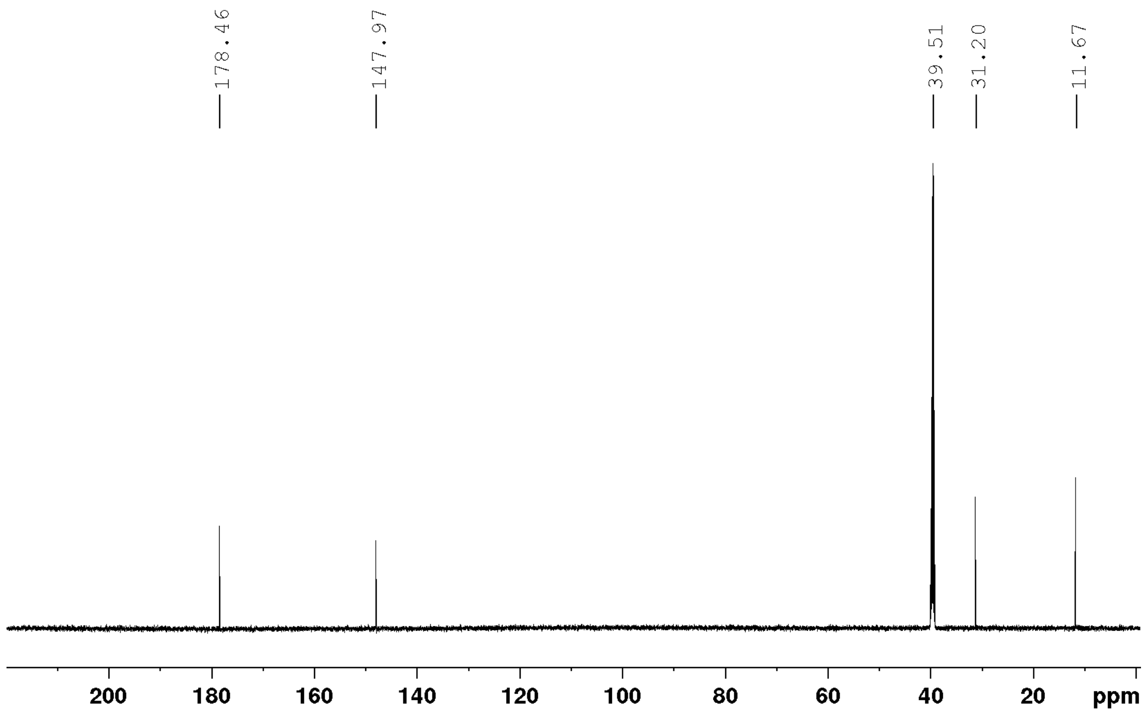


**Supplementary Figure 2. $^{13}C$ NMR Spectrum (125 MHz, DMSO-$d_6$) of $ATSMH_2$.**

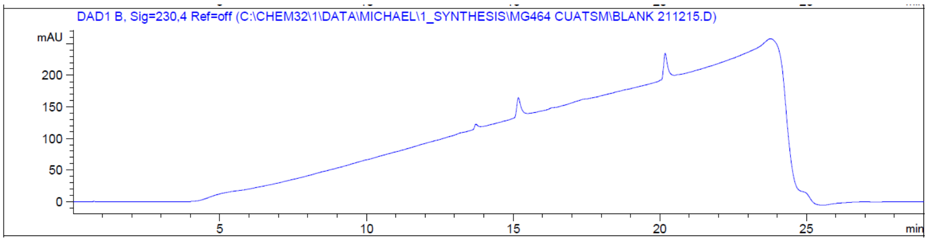


a)

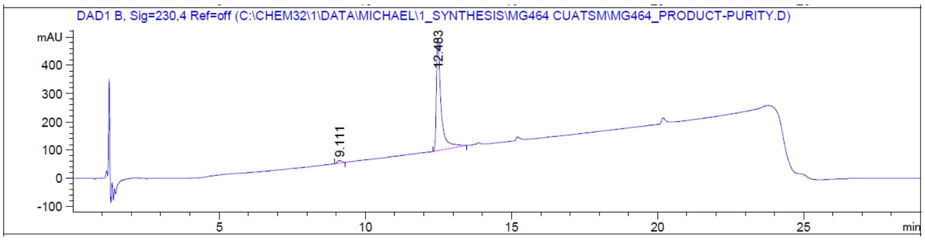


b)

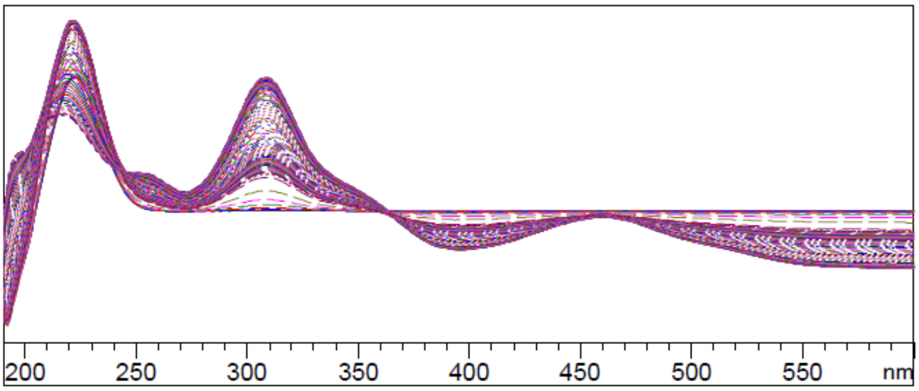


c)

Signal 2: DAD1 B, Sig=230,4 Ref=off

| Peak # | RetTime [min] | Type | Width [min] | Area [mAU*s] | Height [mAU] | Area % |
|---|---|---|---|---|---|---|
| 1 | 9.111 | BB | 0.1304 | 84.95367 | 9.31894 | 1.8851 |
| 2 | 12.483 | BB | 0.1575 | 4421.73096 | 398.25204 | 98.1149 |
| Totals : | | | | 4506.68462 | 407.57099 | |

d)

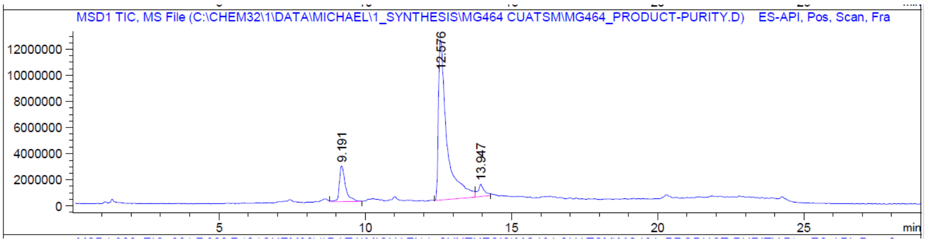


e)

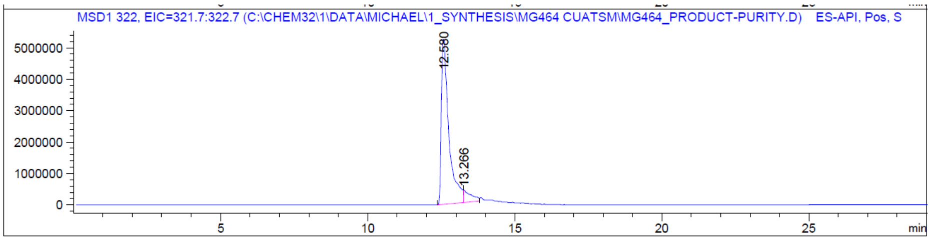


f)

**Supplementary Figure 3. LCMS chromatogram of CuATSM after copper-loading and SPE method development.** UV chromatograms at 230 nm of a) blank (no injection) and b) sample with peaks integrated after subtracting background and solvent front; c) UV spectrum of Cu(II)ATSM; d) purity calculations based on UV absorbances and AUC for product peak and impurities at 230 nm; e) MS chromatogram (ESI+ scan *m/z* 100–1000); and f) Extracted Ion Chromatogram for Cu(II)ATSM ($[M+H]^+$ *m/z* 322.0).

**a)**

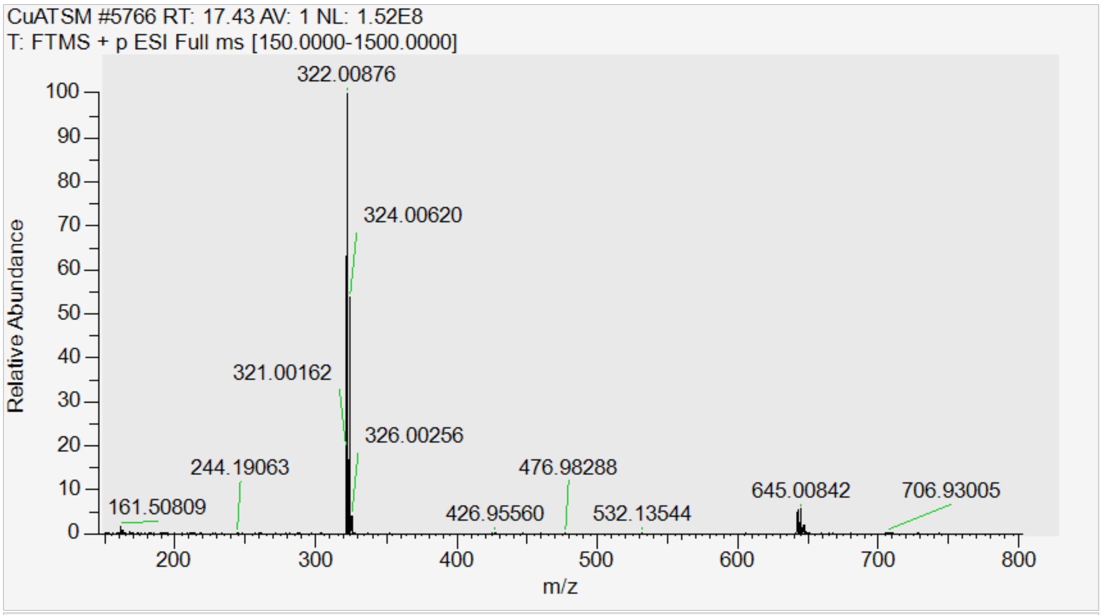


**b)**

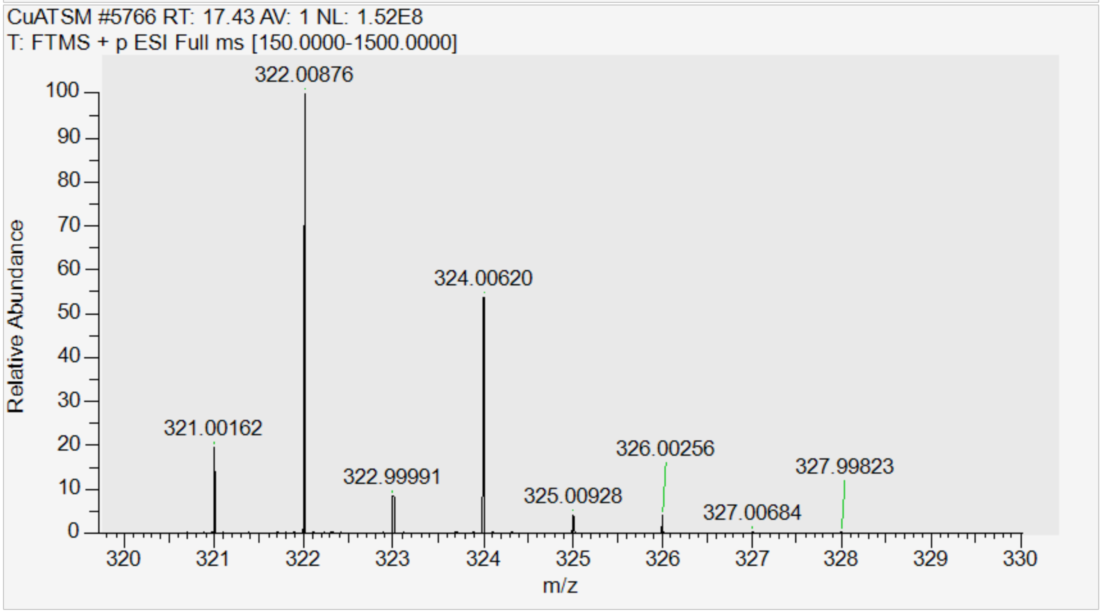


**Supplementary Figure 4. High resolution mass spectrum of CuATSM.** Mass spectrometry found *m/z* 322.00876 (100)/324.0620 (55) for $[M+H^+]^+$), $C_8H_{15}N_6S_2Cu^+$ requires 322.00902/324.00721. a) scan range from *m/z* 150–800; b) enlarged region from m/z 320–330, displaying the distinct isotopic pattern (2:1 for [63]/[65]Cu) for Cu(II)ATSM.

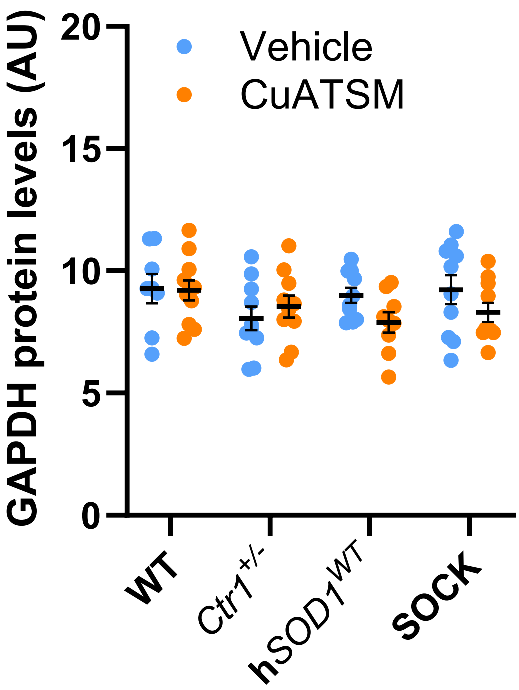


**Supplementary Figure 5. GAPDH protein levels in the midbrain of all mouse strains treated with CuATSM and vehicle.** Measurements were obtained using immunoblotting, representative blots are displayed in **Supplementary Figure 13**. CuATSM treatment did not impact GAPDH levels within any strain, and no significant differences were observed between mouse strains within each treatment group. Full details of statistical tests are presented in **Supplementary Table 2**. Data represents mean ± SEM, *n* = 9-11. Abbreviations: WT, wild-type.

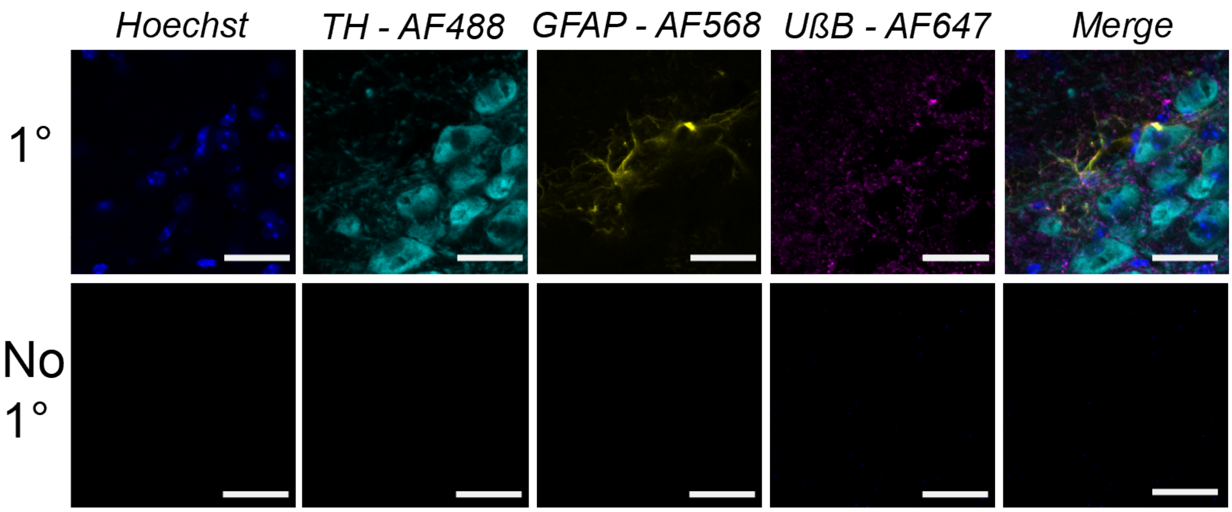


**Supplementary Figure 6. Negative control immunofluorescent staining.** In addition to immunofluorescent staining (primary: 1°) for cell nuclei (Hoechst, blue), tyrosine hydroxylase (TH, cyan), glial acidic fibrillary protein (GFAP, orange) and unfolded beta barrel SOD1 (UβB, magenta), we performed negative control staining (no 1°) by replacing primary antibodies and DAPI with blocking buffer. No fluorescent signals were detected when imaging no 1° staining using the same sequential acquisition settings as those used for imaging 1° staining. Full antibody details are listed in **Supplementary Table 1**. Scale bars represent 30 μm.

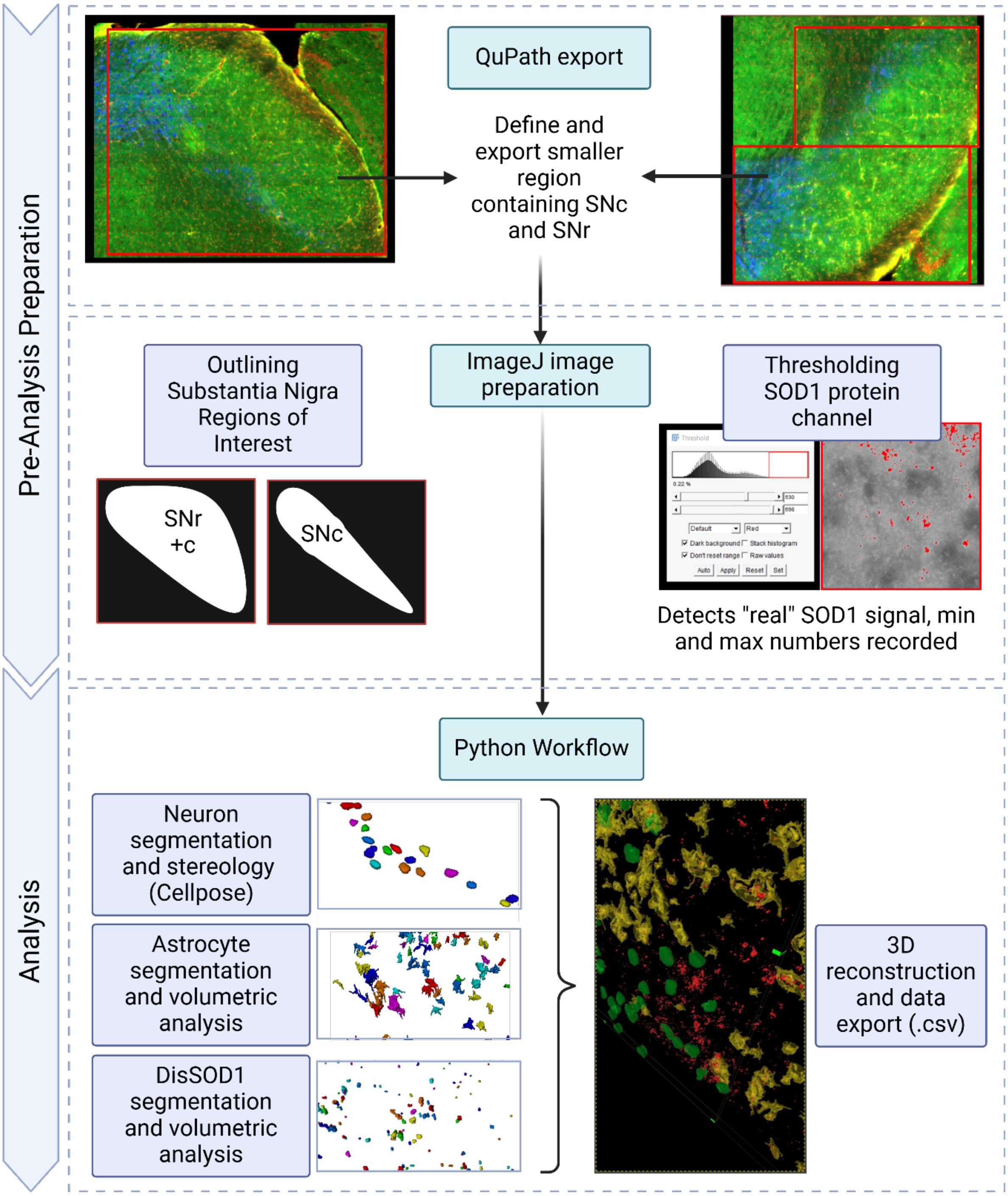


**Supplementary Figure 7. Summary of image processing and analysis workflow used to count dopamine neurons and measure the volume of disSOD1 and astrocytes in the SNc of all mouse strains.** All ImageJ and Python analysis scripts are publicly-available on GitHub (owner: Richard Harwood, repository: Sod1_CuATSM_Image_Analysis).

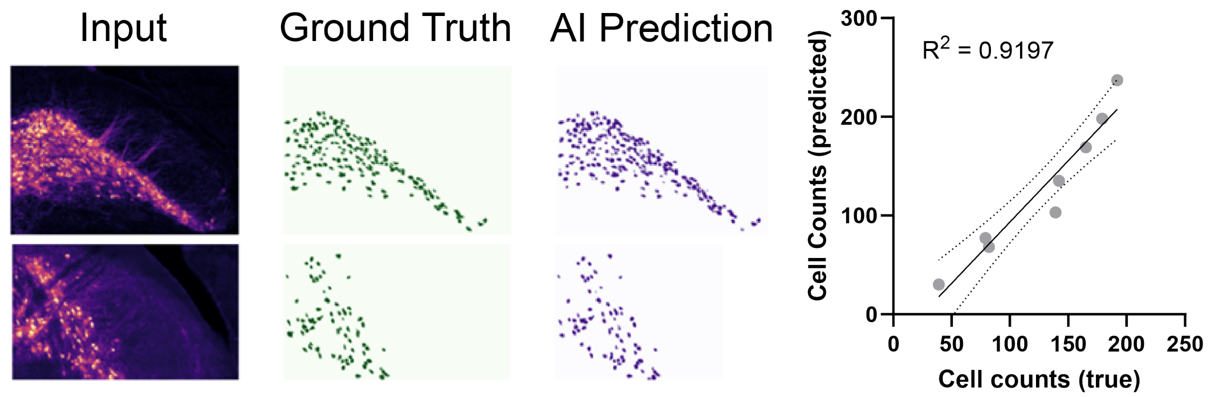


**Supplementary Figure 8. Estimate of the accuracy of re-trained Cellpose deep learning model for segmenting and quantifying SNc dopamine neurons.** Ground truth neuron segmentation (green) was generated via manual segmentation of neurons from input images, whereas AI prediction segmentation (purple) was generated using the retrained Cellpose deep learning model. The difference between ground truth and AI prediction produces the F1 score (0.83). Data points used in linear regression analysis represent counts obtained for individual z stacks from different mice.

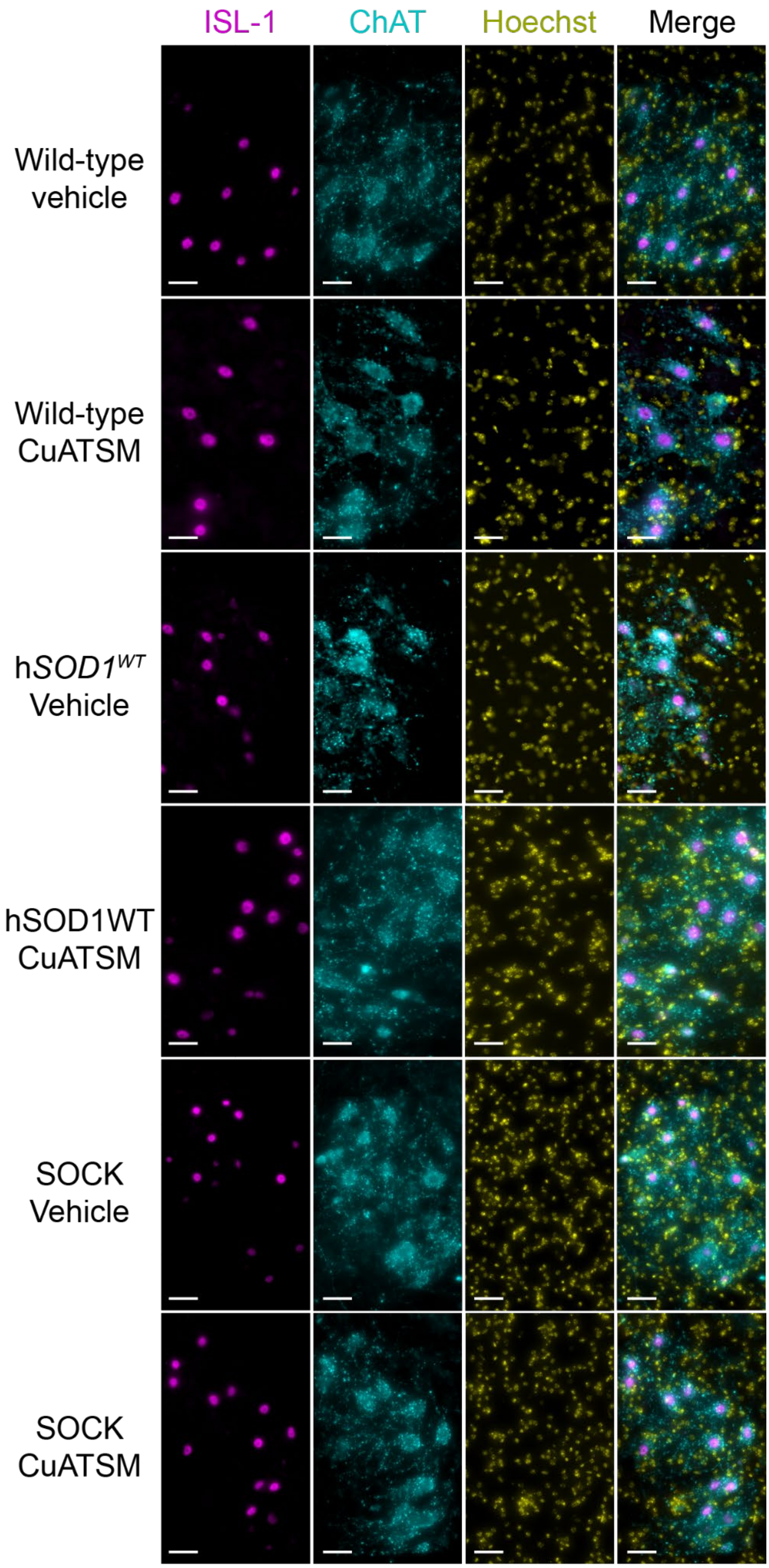


**Supplementary Figure 9. Immunofluorescent staining for spinal motor neurons in all four mouse genotypes following treatment.** Antibody details are listed in **Supplementary Table 1**.

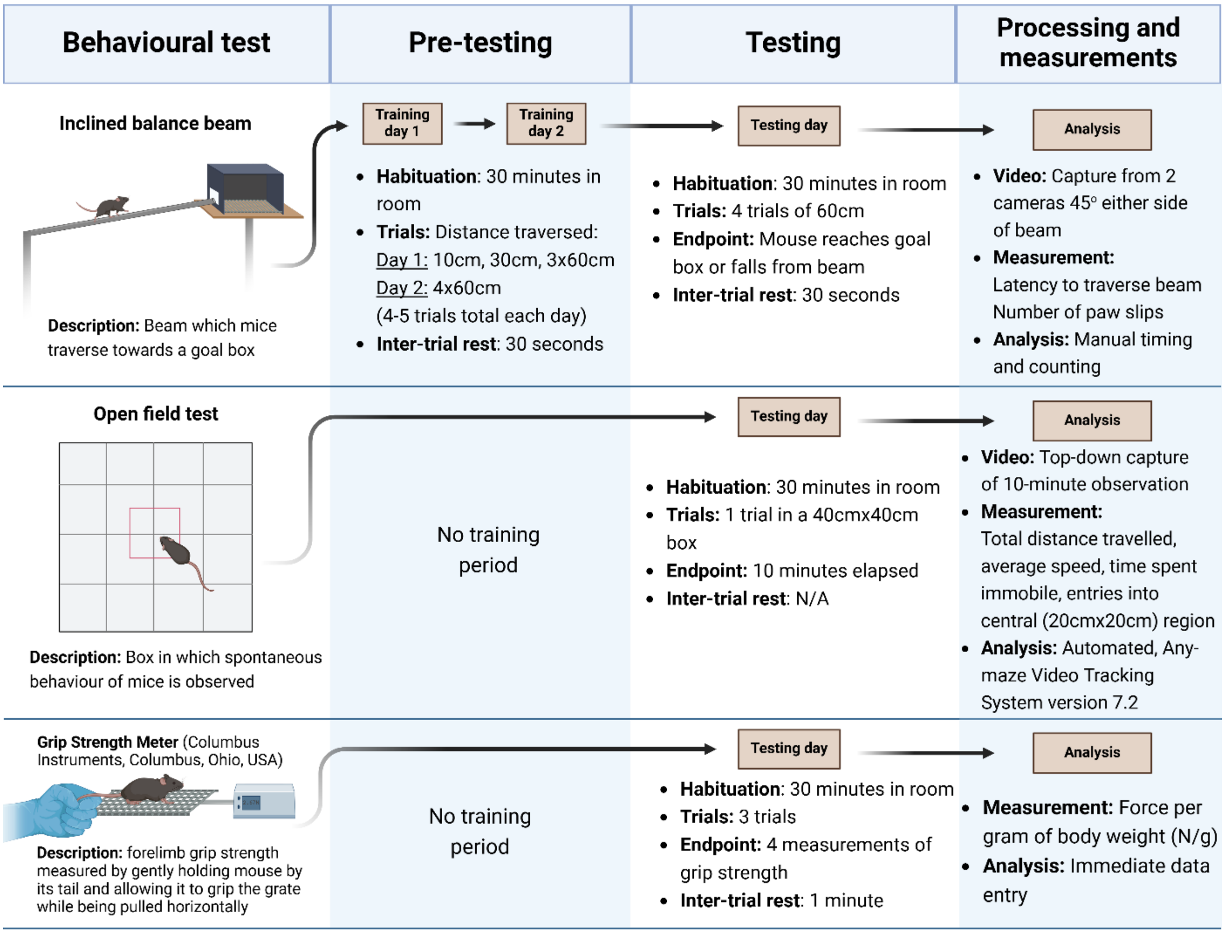


**Supplementary Figure 10. Summary of behavioural test procedures performed on all mice.** Tests were performed at 5.5 months-of-age immediately prior to animal culling. N=Newtons, rpm=revolutions per minute.

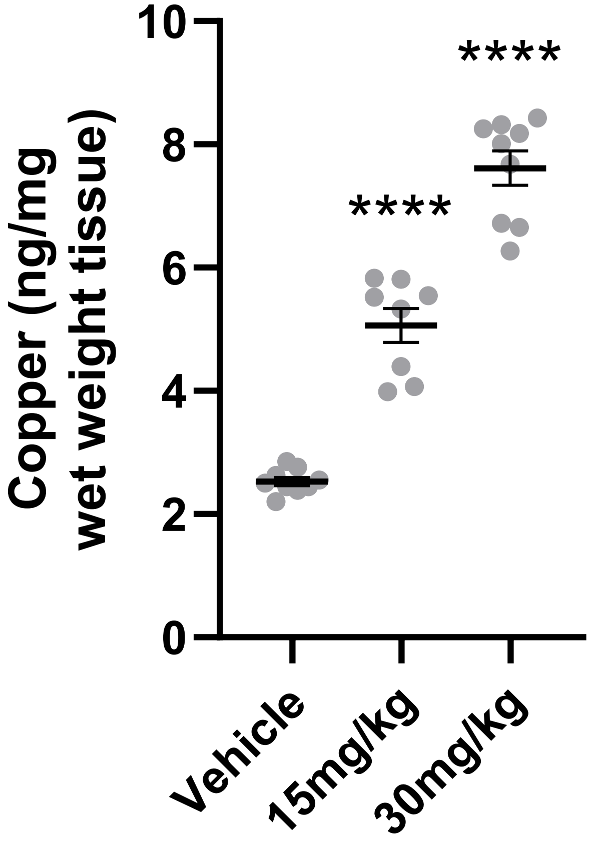


**Supplementary Figure 11. Pilot study to determine CuATSM dose.** Midbrain copper levels were quantified using inductively-coupled plasma mass spectrometry in wild-type C57BL/6 mice treated with either 15 or 30mg/kg CuATSM, or vehicle, for three weeks. Data represent mean ± SEM ($n$ = 8-9/treatment), with full details of statistical tests presented in **Supplementary Table 2**. ****$p < 0.0001$.

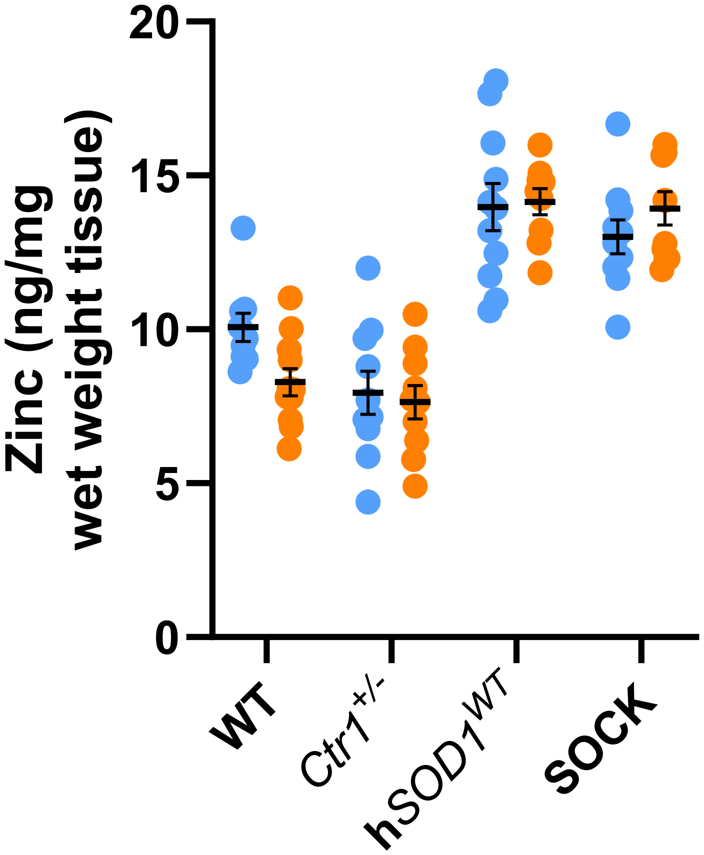


**Supplementary Figure 12. Zinc levels in the midbrain of all four mouse strains treated with vehicle or CuATSM.** Quantification was performed using inductively-coupled plasma mass spectrometry. Data represent mean ± SEM (*n* = 10/genotype/treatment), with full details of statistical tests presented in **Supplementary Table 2**.

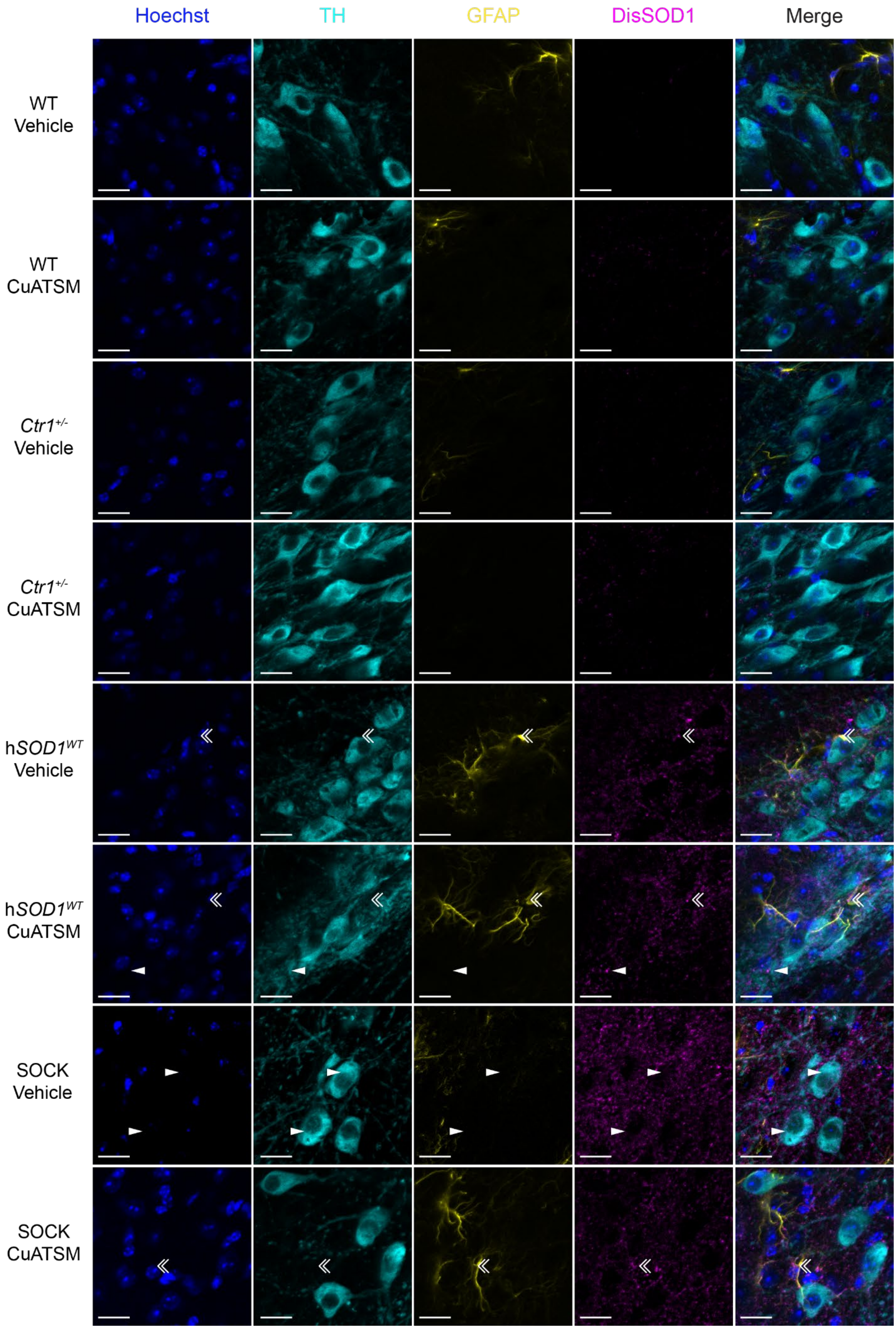
Hoechst
TH
GFAP
DisSOD1
Merge
WT Vehicle
WT CuATSM
Ctr1+/- Vehicle
Ctr1+/- CuATSM
hSOD1WT Vehicle
hSOD1WT CuATSM
SOCK Vehicle
SOCK CuATSM

**Supplementary Figure 13. Immunofluorescent characterization of disSOD1 pathology in the SNc of all four mouse strains treated with vehicle or CuATSM.** Tyrosine hydroxylase (TH) was used to identify dopaminergic neurons, glial fibrillary acidic protein (GFAP) was used to identify astrocytes, while the unfolded β-barrel conformation-specific SOD1 antibody was used to delineate disSOD1 pathology. Antibody details are presented in **Supplementary Table 1**. White arrowheads identify disSOD1 deposits within dopamine neurons, while double white arrowheads delineate those inside astrocytes. Scale bars represent 20 µm.

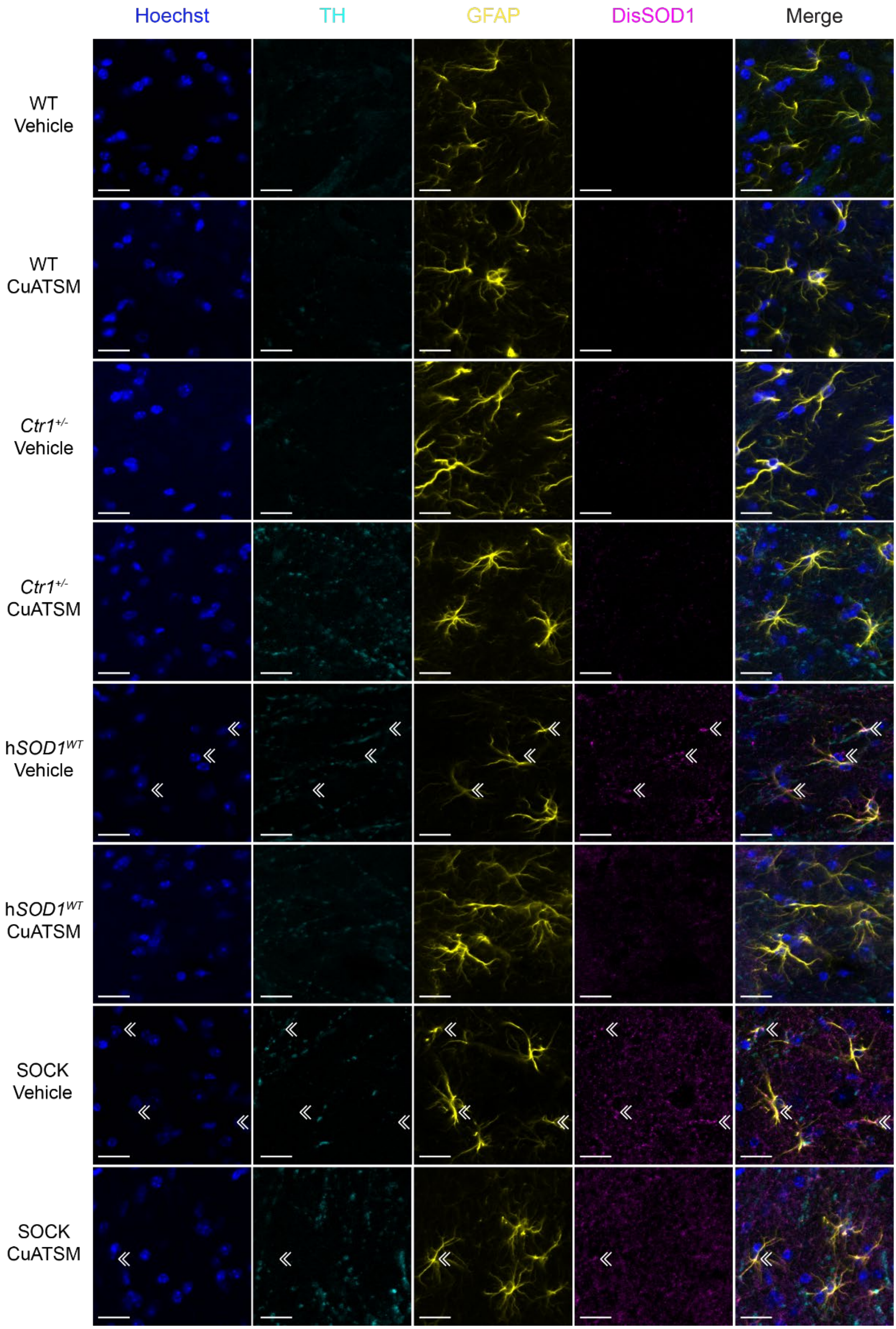
Hoechst
TH
GFAP
DisSOD1
Merge
WT
Vehicle
WT
CuATSM
Ctr1+/-
Vehicle
Ctr1+/-
CuATSM
hSOD1WT
Vehicle
hSOD1WT
CuATSM
SOCK
Vehicle
SOCK
CuATSM

**Supplementary Figure 14. Immunofluorescent characterization of disSOD1 pathology in the SNr of all four mouse strains treated with vehicle or CuATSM.** Tyrosine hydroxylase (TH) was used to identify dopamine neuron processes, glial fibrillary acidic protein (GFAP) was used to identify astrocytes, while the unfolded β-barrel conformation-specific SOD1 antibody was used to delineate disSOD1 pathology. Antibody details are presented in **Supplementary Table 1**. White arrowheads identify disSOD1 deposits within dopamine neurons, while double white arrowheads delineate those inside astrocytes. Scale bars represent 20 μm.

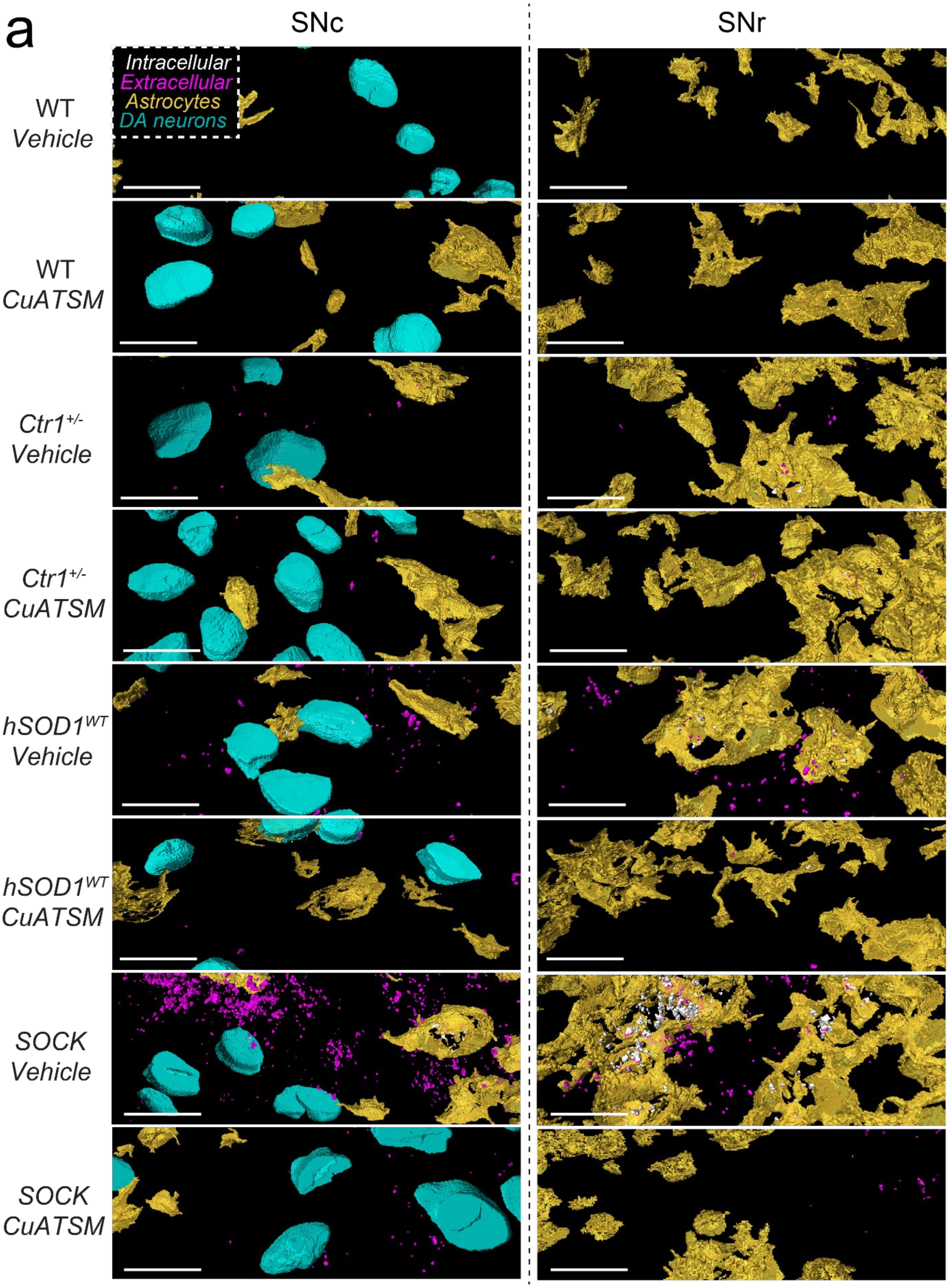


**Supplementary Figure 15. Three dimensional reconstructions of disSOD1 pathology in the SNc and SNr of all mouse strains treated with vehicle and CuATSM.** Tyrosine hydroxylase (TH)-positive neuronal soma are highlighted in cyan, astrocytes in yellow, while aggregates residing within and outside of these cell types are highlighted in white and magenta, respectively. Scale bars represent 20µm.

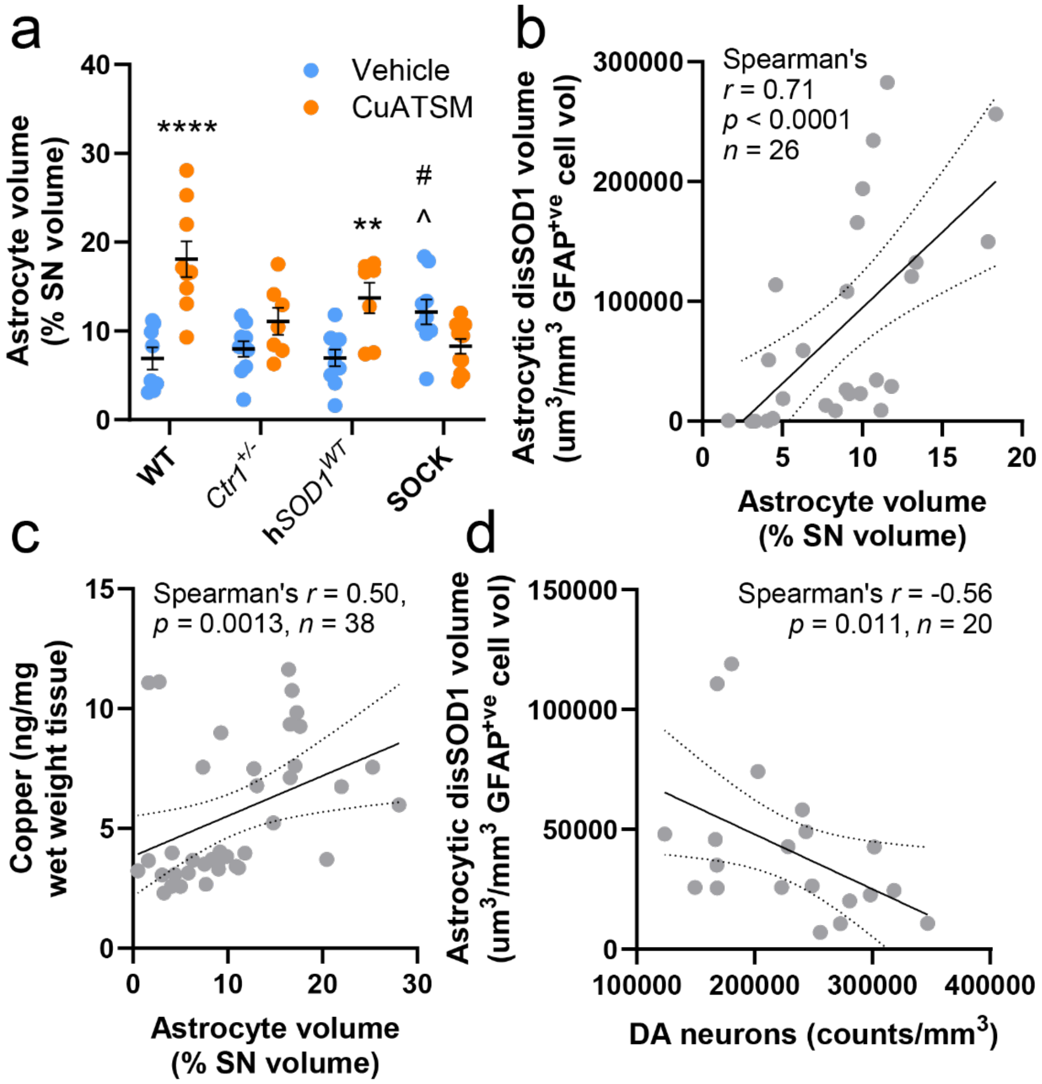


**Supplementary Figure 16. Astrocyte volume in the SN of all mouse strains following vehicle and CuATSM treatment. a.** Quantification was performed using volumetric analysis of three-dimensional reconstructions of GFAP immunostaining in the SN, demonstrating an increase in astrocyte volume in vehicle-treated SOCK mice compared with wild-type and h*SOD1*$^{WT}$ mice, as well as in wild-type and h*SOD1*$^{WT}$ mice following CuATSM treatment. Data represent mean ± SEM (*n* = 9-11/genotype/treatment), with full details of statistical tests presented in **Supplementary Table 2**. Comparisons marked with an asterisk (*) denote those made between vehicle- and CuATSM-treated mice of the same genotype, those marked with an arrowhead (^) represent those made to vehicle-treated wild-type mice, while those marked with a hashtag (#) signify those made to vehicle-treated h*SOD1*$^{WT}$ mice. # $p < 0.05$, ****$p < 0.0001$, ** $p < 0.01$, ^$p < 0.05$. **b.** Astrocyte volume in vehicle-treated wild-type, h*SOD1*$^{WT}$ and SOCK mice was correlated with the volume of disSOD1. **c.** Astrocyte volume in the midbrains of vehicle- and CuATSM-treated wild-type and h*SOD1*$^{WT}$ mice was correlated with tissue copper content (**Fig. 1e**). **d.** Astrocytic disSOD1 volume was negatively correlated with dopamine (DA) neuron density in the SNc of vehicle- and CuATSM-treated SOCK mice. Details of statistical tests for **b** - **d** are presented in each panel.

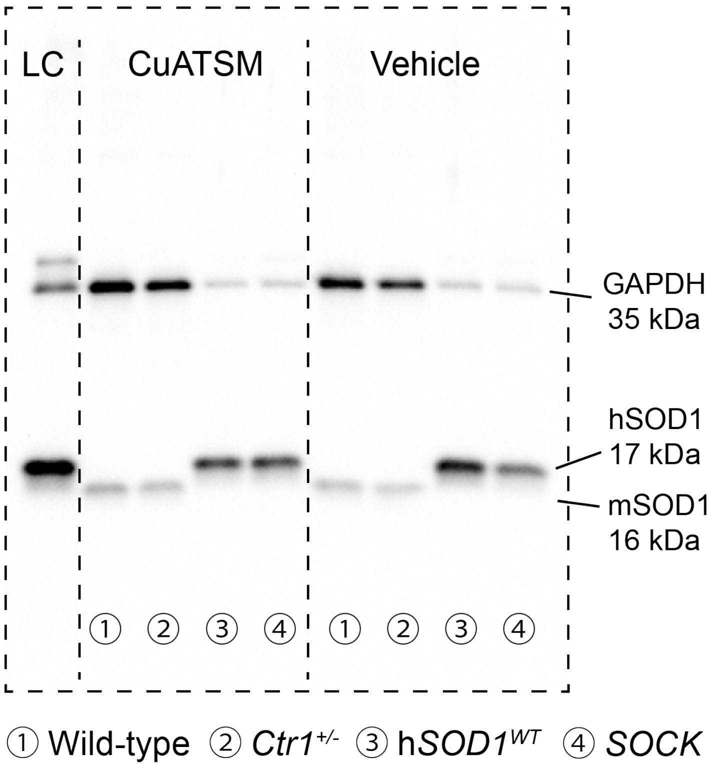


**Supplementary Figure 17. Representative SOD1 and GAPDH immunoblots from midbrain tissues of all four mouse genotypes.** A common loading control (LC) made from pooled midbrain, cortex and liver extracts was included on all gels to enable standardization and estimation of protein expression differences between these groups. Blots were probed simultaneously for SOD1 (human isoform; 16 kDa, mouse isoform; 17 kDa) and GAPDH (two bands; 35 and 36 kDa) given their distinct molecular weights. Full antibody details are listed in **Supplementary Table 1**.

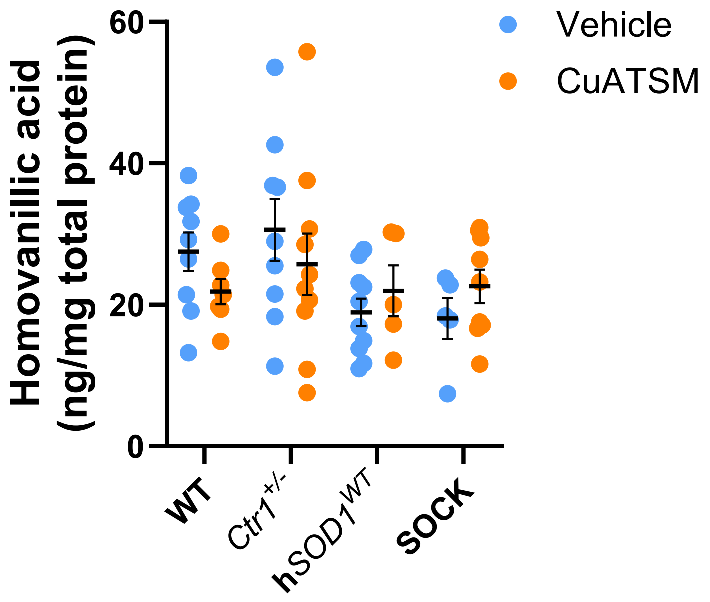


**Supplementary Figure 18. Homovanillic acid levels in the striatum of all four mouse strains treated with vehicle or CuATSM.** No differences in striatal homovanillic acid levels were identified between mouse strains or treatment groups using high performance liquid chromatography. Data represent mean ± SEM (*n* = 9-11/genotype/treatment), with full details of statistical tests presented in **Supplementary Table 2**.

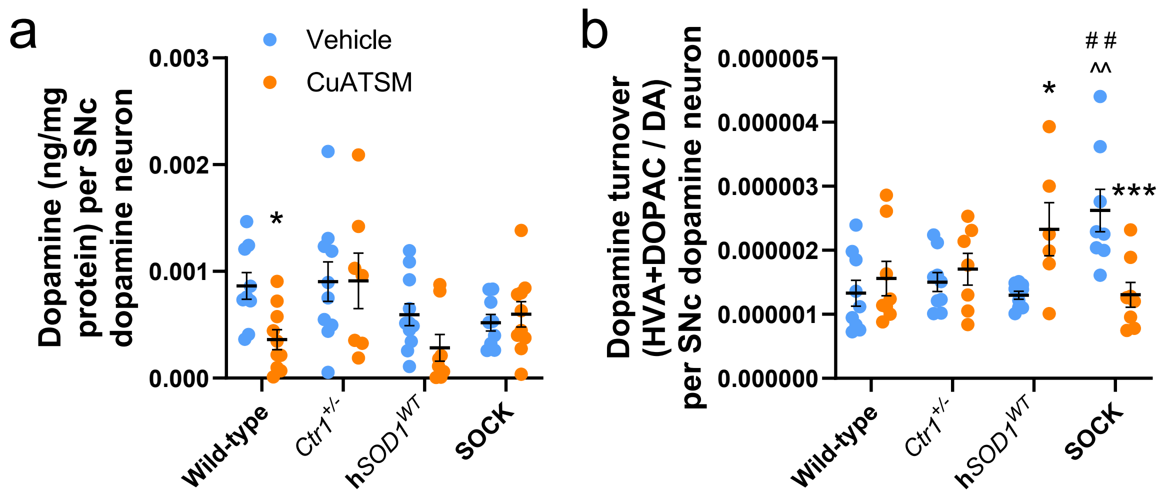


**Supplementary Figure 19. Striatal dopamine levels and turnover normalized to SNc dopamine neuron density in all four mouse strains treated with vehicle or CuATSM. a.** Normalized dopamine levels only differed in wild-type mice between vehicle- and CuATSM-treated mice. **b.** Dopamine turnover was significantly increased in hSOD1WT mice following CuATSM treatment but was decreased in CuATSM-treated SOCK mice. Comparisons marked with an asterisk (*) denote those made between vehicle- and CuATSM-treated mice of the same genotype, those marked with an arrowhead (^) specify those made to vehicle-treated wild-type mice, while those marked with a hashtag (#) denote those made to vehicle-treated h$SOD1^{WT}$ mice. *$p < 0.05$, ***$p < 0.001$, ^^$p < 0.01$, # # $p < 0.01$. Data represent mean ± SEM ($n$ = 9-11/genotype/treatment), with full details of statistical tests presented in **Supplementary Table 2**.
